# Targeting *MALAT1* Augments Sensitivity to PARP Inhibition by Impairing Homologous Recombination in Prostate Cancer

**DOI:** 10.1101/2022.06.01.494272

**Authors:** Anjali Yadav, Tanay Biswas, Ayush Praveen, Promit Ganguly, Ayushi Verma, Dipak Dutta, Bushra Ateeq

## Abstract

Poly(ADP-ribose) polymerase inhibitors (PARPi) have emerged as the most promising targeted therapeutic intervention for the treatment of metastatic castrate-resistant prostate cancer (mCRPC). However, the clinical utility of PARPi has been limited to a subset of patients who harbor aberrations in the homologous recombination (HR) pathway. Here, we report that targeting *MALAT1,* an oncogenic lncRNA, known to be elevated in advanced-stage prostate cancer (PCa) demonstrates contextual synthetic lethality with PARPi. We show that *MALAT1* silencing reprograms the HR transcriptome, contriving BRCAness-like phenotype, thus enhancing sensitivity towards PARPi. Moreover, transcriptome profiles of mCRPC patients exhibit convergence between expression of *MALAT1*, HR pathway, and neuroendocrine markers. Mechanistically, we show that targeting *MALAT1* leads to a decrease in EZH2, a member of polycomb repressor complex-2 (PRC2), which in turn upregulates the expression of RE1 Silencing Transcription Factor (REST), a key repressor of neuroendocrine differentiation. Overall, we showed that *MALAT1* plays a pivotal role in maintaining genomic integrity, thereby promoting disease progression. Conclusively, our findings suggest that inhibiting *MALAT1* confers PARPi sensitization in patient’s resistant to anti-androgens and conventional chemotherapeutics.

**Graphical abstract:**
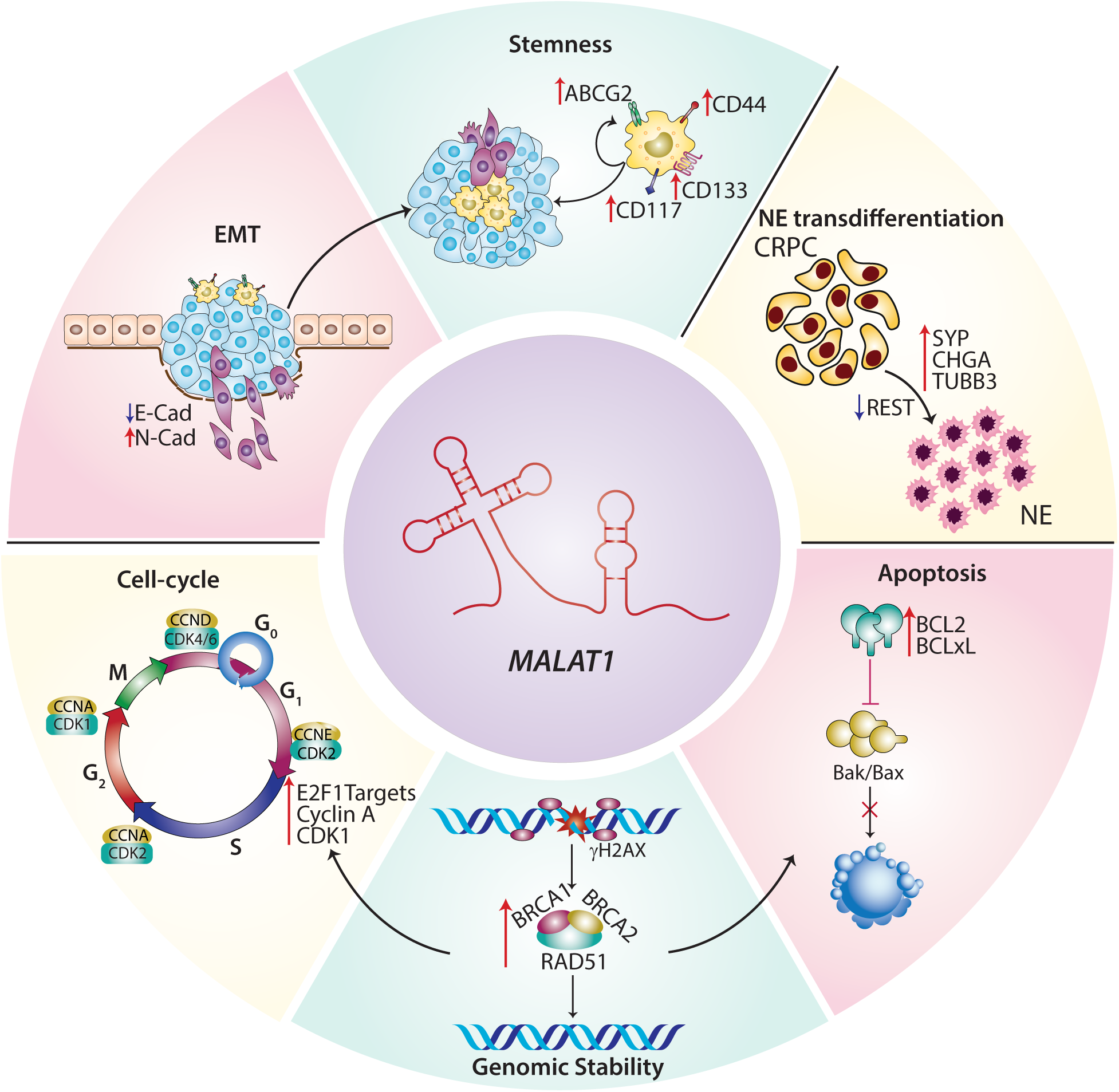
The oncogenic lncRNA *MALAT1* exhibits functional pleiotropy in PCa and promotes neuroendocrine differentiation. *MALAT1* fosters PCa progression by modulating several hallmark oncogenic properties, such as malignant transformation, enhanced migratory capabilities, stemness, and ultimately contributes to drug resistance. *MALAT1* enhances the transcriptional regulation of genes associated with homologous recombination thereby having a profound impact on the genome integrity in metastatic prostate cancer. Over the course of disease progression, it also promotes neuroendocrine trans-differentiation by depleting the levels of REST, the key repressor for NE transdifferentiation in prostate cancer.

## INTRODUCTION

Androgen deprivation is the most preferred systemic therapy for the treatment of advanced-stage prostate cancer (PCa) (1), which initially mitigates the disease, nonetheless, most patients inevitably develop castration-resistant prostate cancer (CRPC) (2, 3). However, CRPC tumors restore androgen signaling via several alternative mechanisms such as somatic mutations in androgen receptor (AR), expression of constitutively active AR splice variant (AR-V7), intratumoral androgen synthesis, and mutations in coactivators and/or corepressors (4). Hence, second-generation androgen signaling inhibitors (such as apalutamide, darolutamide) either alone or in combination with taxanes (e.g., docetaxel or cabazitaxel) are used for the treatment of CRPC patients (5). Despite the fact that these medications substantially alleviate the symptoms and prolong the patient’s survival, none of these are effective in the long term (6–8). Furthermore, a subset of aggressive CRPC patients often develops a more aggressive phenotype, known as neuroendocrine prostate cancer (NEPC) marked by upregulation of synaptophysin (SYP), enolase 2 (ENO2), chromogranin A (CHGA) and SPINK1 (9–11). NEPC primarily originates from a selective outgrowth of AR negative subpopulation of CRPC cells, and deregulation of several transcription factors, namely MYCN, EZH2, REST, Aurora kinase A, and B (AURKA/B), BRN2, PEG10, SRRM4, SOX2 (12–17). Recently, Ramnarine et. al. showed that long noncoding RNAs (lncRNAs), such as *H19*, *LINC00617*, and *SSTR5-AS1* are upregulated in NEPC (18), but their biological significance is yet to be investigated.

Genetic alterations in the DNA damage response (DDR) genes including loss of function mutations in *BRCA1/2* has been observed in ∼20-30% of advanced stage PCa patients (19–21). It has been shown that tumor cells harboring mutations in DDR genes are highly susceptible to PARP inhibitors due to synthetic lethality (22–24). The clinical utility of PARPi is limited to patients harboring HR-deficient tumors, nonetheless, these patients often develop reversion mutations in the HR genes and become non-responsive (25, 26). Mounting evidence suggests that chemotherapeutic drugs that potentiate DNA damage or induce HR deficiency can increase sensitivity to PARP inhibitors. Although, these combinatorial approaches increase the duration of response in HR-deficient tumors, and expand the utility of PARPi for HR-proficient cancers, but are not well tolerated as patients often develop debilitating side effects such as myelosuppression (27, 28). This highlights the necessity to discover novel and targetable molecular modulator(s) that could facilitate multistep targeting of the HR pathway to induce “BRCAness” like physiological state in cancer.

Here, we deciphered a molecular network between lncRNA *MALAT1*, HR pathway, AR signaling, and neuroendocrine phenotype in mCRPC patients. We showed that *MALAT1* modulates DNA repair pathways and maintains genome integrity in prostate cancer. Additionally, higher *MALAT1* levels facilitate NE transdifferentiation by downregulating the expression of a crucial neuronal repressor, REST by interacting with EZH2. Collectively, our findings point towards a novel therapeutic approach wherein targeting *MALAT1* augments sensitivity to PARP inhibition by inducing HR deficiency in prostate cancer.

## RESULTS

### *MALAT1* is highly expressed in metastatic prostate cancer and associates with poor prognosis

To identify key genes associated with advanced stage PCa, we performed differential gene expression analysis using three publicly available microarray datasets comprising expression profiles of PCa patients, namely GSE35988 (29), GSE6919 (30), and GSE6752 (31). A total of 163 transcripts consisting of 161 genes, a pseudogene *RPL32P3,* and a long non-coding RNA (lncRNA), *MALAT1* were found to be significantly elevated in metastatic PCa patients compared to localized cases (**Figure 1A; Supplemental Table S1**). We observed *MALAT1* was highly upregulated in metastatic cases (∼1.5–4 times) compared to localized PCa patients (**Figure 1, B-D)**. Analysis of transcriptome data from The Cancer Genome Atlas Prostate Adenocarcinoma (TCGA-PRAD) cohort (32) revealed that *MALAT1* (n=499) expression positively correlates with several clinicopathological parameters such as Gleason score and node status (**Supplemental Figure S1A**). Moreover, elevated levels of *MALAT1* were observed in patients with progressive disease (PD) compared to those with complete response (CR) after primary therapy (**Supplemental Figure S1A**), indicating its significance in predicting response to chemotherapy. Besides, higher *MALAT1* was also noted in the patients who suffered from biochemical recurrence (BCR; **Supplemental Figure S1B**). To further validate, we next performed a Kaplan-Meier survival analysis by stratifying the TCGA-PRAD patients by their median expression into high (⩾median, n=248) and low (⩽median, n=250) groups. Intriguingly, patients with elevated *MALAT1* levels showed a higher probability of BCR compared to the *MALAT1*-low group (*p*=0.016; **Figure 1E**). In addition, *MALAT1*-high patients in the TCGA-PRAD cohort showed higher chances of relapse compared to the *MALAT1*-low group (*p*=0.0013; **Figure 1F**), suggesting that *MALAT1* levels could also predict the likelihood of disease recurrence.

**Figure 1:**
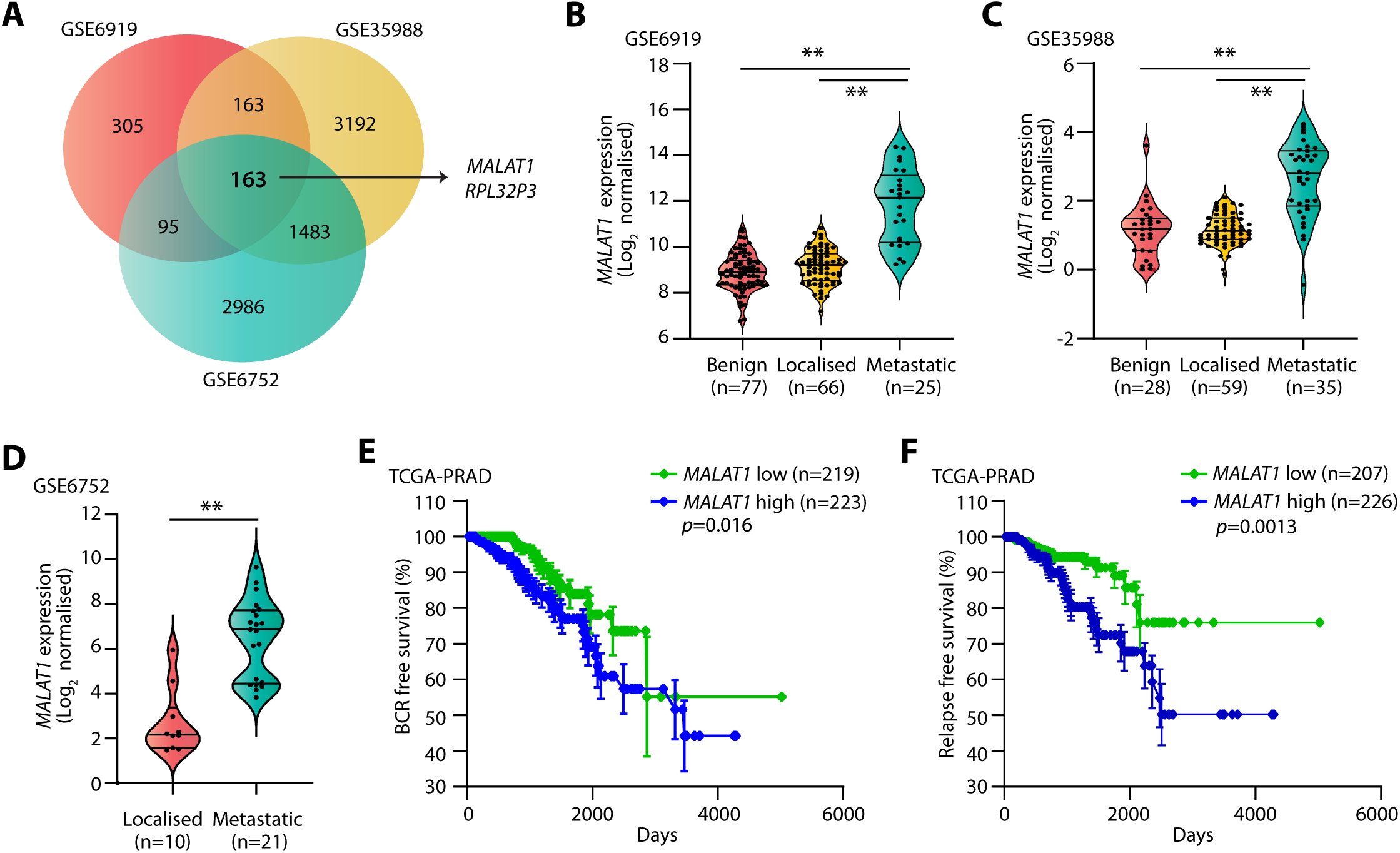
High *MALAT1* expression associates with poor prostate cancer prognosis. **A.** Venn diagram displaying genes upregulated in metastatic PCa patients compared to localized cases in three publicly available Gene Expression Omnibus (GEO) datasets namely, GSE35988, GSE6919, and GSE6752. **B.** Dot plot with superimposed violin plot showing *MALAT1* expression in benign, primary, and metastatic PCa patients in the GSE6919 dataset. *MALAT1* transcript is reported as log_2_ median-centred ratio. **C.** Same as B, except for the GSE35988 dataset. **D.** Same as B, except GSE6752 dataset. **E.** Kaplan-Meier curves for biochemical recurrence-free survival in TCGA-PRAD dataset categorized as “*MALAT1*-High” (n =248) and “*MALAT1*-Low” (n = 250) groups based on the median expression of *MALAT1.* Blue line represents patients with higher expression of *MALAT1* whereas the green line represents lower. The *p*-values were computed by log-rank test. **F.** Same as D, except relapse-free survival for TCGA-PRAD dataset. Data represent mean ± SEM. For panel B-E one-way ANOVA with Dunnett’s multiple comparisons, a posthoc test was applied and for panel F two-tailed unpaired Student’s *t-*test was applied. ∗*p* ≤ 0.05 and ∗∗*p* ≤ 0.001.

RNA-seq data analyses from the cancer cell line encyclopedia (CCLE) revealed high expression of *MALAT1* in multiple human cancer cell lines, wherein the highest expression was noted in PCa (**Supplemental Figure S1C**). Similarly, our quantitative PCR (qPCR) data with multiple PCa cell lines (PC3, DU145, LNCaP, 22RV1, and VCaP) also exhibited higher expression of *MALAT1* compared to PNT2, an immortalized non-tumorigenic normal prostate epithelial cell line (**Supplemental Figure S1D**). Taken together, these results indicate that *MALAT1* positively correlates with the clinicopathological features of aggressive PCa, and may serve as a promising prognostic marker.

### *MALAT1* promotes EMT, stemness, and chemoresistance in prostate cancer

*MALAT1* emerged as a metastasis-promoting lncRNA in multiple malignancies and serves as a predictor in assessing response to cancer therapies (33–35). As indicated in Figure 1, tumors with elevated levels of *MALAT1* show a higher propensity for lymph node metastasis, disease recurrence, and therapeutic failure. Therefore, we examined the association of *MALAT1* with molecular factors allied with epithelial-to-mesenchymal transition (EMT) in the TCGA-PRAD cohort. Intriguingly, *MALAT1-*low patients showed increased levels of archetypal epithelial markers, such as *CHD1, SPINT2, EPCAM, TJP1, CLDN7,* and *OCL* compared to *MALAT1*-high patients (**Supplemental Figure S1E).** Likewise, *MALAT1-*high patients showed elevated levels of mesenchymal markers, such as *CTGF, KRT5, FOXC1, EMP3, SNAI1,* and *FOXC2* (**Supplemental Figure S1E).** To further investigate the functional significance of *MALAT1* in invasion and metastasis, we generated stable *MALAT1*-silenced (sh*MALAT1*) and scrambled control (SCRM) 22RV1 and LNCaP cells using lentivirus-based short-hairpin RNAs (**Supplemental Figure S1F)**. Characterization of these stable cells revealed a robust increase in the epithelial marker, E-cadherin, accompanied with a significant decrease in mesenchymal marker (N-cadherin) in sh*MALAT1* cells compared to respective SCRM control, emphasizing its importance in EMT (**Figure 2A**). In accord with this, *MALAT1* depletion dramatically reduced 3D cell migration in 22RV1 (∼60%) and LNCaP (∼70%) cells compared to respective SCRM controls (**Figure 2B**), suggesting that *MALAT1* is required for migration in PCa cells.

**Figure 2:**
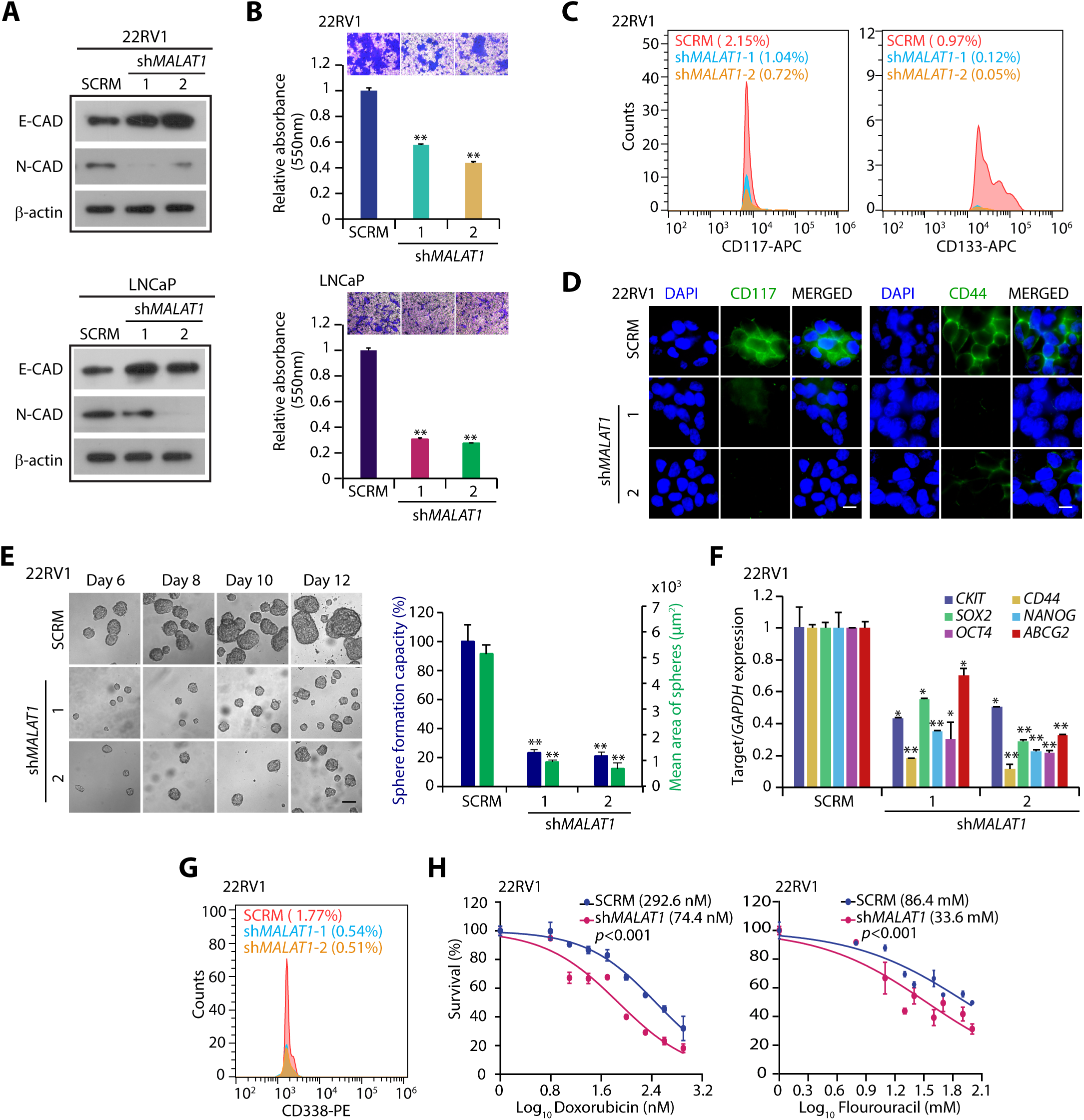
*MALAT1* promotes EMT, stemness, and chemoresistance in prostate cancer. **A.** Immunoblots showing the expression of EMT markers in sh*MALAT1* and SCRM PCa cells. β-actin was used as an internal control. **B.** Boyden chamber Matrigel migration assay using same cells as in **A**. Representative fields with the migrated cells are shown in the inset. Bar plot depicts alteration in migratory potential of the PCa cells upon *MALAT1* knockdown. **C.** Flow cytometry analysis showing expression of CD117 (c-KIT) and CD133 in 22RV1-sh*MALAT1* and SCRM control cells. **D.** Immunofluorescence images displaying the expression of CD117 and CD44 and in the same cells as in **C**. **E.** Representative phase-contrast images for the prostatospheres formed using 22RV1-sh*MALAT1* and SCRM control cells on the indicated days. Right panel: Bar plot representing percentage sphere formation efficiency and the mean area of the prostatospheres; Scale bar represents 100 μm. **F.** QPCR depicting the expression of stem cell markers in the prostatospheres derived from the same cells as in **E**. Expression level for each gene was normalized to *GAPDH*. **G.** Flow cytometry analysis showing expression of ATP-binding cassette superfamily G member 2 (CD338/ABCG2) using the same cells as in **C.** **H.** Cell cytotoxicity assay using chemotherapeutic drugs namely, doxorubicin and 5-fluorouracil using the same cells as in C IC_50_ values were calculated by generating a dose-response curve using Graph-pad Prism software. Experiments were performed with n=3 biologically independent samples; data represents mean ± SEM. For panels, B, E, and F one-way ANOVA with Dunnett’s multiple-comparisons posthoc test was applied. ∗*p* ≤ 0.05 and ∗∗*p* ≤ 0.001.

Metastatic PCa frequently acquires cancer stem cell properties, which result in self-renewal and chemoresistance (36, 37), we next examined the TCGA-PRAD cohort for any association of *MALAT1* with self-renewal factors. Interestingly, *MALAT1-*high patients showed increased levels of several key stemness factors, such as *Octamer-binding transcription factor 4 (OCT4), Homeobox protein NANOG, Krüppel-like factor 4 (KLF4), ATP binding cassette subfamily G member 2 (ABCG2), Sex Determining Region Y-Box 2/9 (SOX2* and *SOX9)* compared to *MALAT1*-low patients (**Supplemental Figure S1G**), indicating positive association of *MALAT1* with stemness. To confirm this, we examined the expression of surface proteins associated with cancer stem cells (CSCs), namely CD117 (c-KIT), a tyrosine kinase receptor; CD331, human fibroblast growth factor receptor 1, and CD44, a cell-surface glycoprotein in 22RV1-sh*MALAT1* and -SCRM cells. Interestingly, sh*MALAT1* cells displayed a marked reduction in the expression of CD117 (∼50-70%), CD133 (∼90%), and CD44 (∼80%) compared to SCRM control (**Figure 2, C and D**). Since self-renewal of tumor cells is the key hallmark of stemness, we next examined the prostatosphere formation ability using 22RV1-sh*MALAT1* and -SCRM cells (37). As anticipated, *MALAT1* depletion abrogated the prostatosphere forming ability of 22RV1 cells as well as their expansion in subsequent serial propagations (**Figure 2E**). Moreover, a significant decrease in the size of prostatospheres was also noted in *MALAT1* deficient cells (**Figure 2E**). Molecular characterization of 22RV1-sh*MALAT1* prostatospheres revealed a marked reduction in the expression of pluripotency genes, namely *C-KIT, OCT-4*, *NANOG*, *CD44*, *SOX2,* and ABCG2 (**Figure 2F**), Likewise, a marked reduction in the expression of CD338 (ABCG2), an ATP-binding cassette transporter was observed in 22RV1 sh*MALAT1* cells as compared to 22RV1-SCRM (**Figure 2G**), suggesting that *MALAT1* modulates stemness in PCa cells.

In addition to stem cell maintenance, several pluripotency factors such as CD117 and ABCG2 have been shown to confer resistance to chemotherapeutic agents. In line with this, Liu et. al. reported that a subpopulation of 22RV1 cells that overexpressed CD117 and ABCG2 exhibited multi-drug resistance (38). Hence, we next investigated the susceptibility of *MALAT1* silenced cells to chemotherapeutic drugs, namely doxorubicin and 5-fluorouracil (5-FU). Interestingly, 22RV1-sh*MALAT1* cells exhibited enhanced sensitivity to chemotherapeutic drugs compared with 22RV1-SCRM (**Figure 2H**). These findings thus provide compelling evidence that downregulation of *MALAT1* effectively suppresses EMT and stemness, and confers sensitivity to chemotherapeutic agents.

### *MALAT1* regulates DNA repair and maintains genome integrity in metastatic prostate cancer

To further delve into the molecular mechanism(s) underlying *MALAT1* mediated carcinogenesis in the prostate, we analyzed the transcriptome profiles of *MALAT1* silenced and control LNCaP-abl cells, retrieved from a publicly available dataset, GSE72534 (39). Our analysis revealed 4139 differentially expressed transcripts with log_2_ FC (FPKM⩽-0.6 or ⩾0.6) that includes ∼1,986 up- and ∼2,153 down-regulated genes in *MALAT1* silenced LNCaP-abl cells compared to the control cells. Functional annotation of the differentially downregulated genes using DAVID (Database for Annotation, Visualization, and Integrated Discovery (40) demonstrates several pathways associated with DNA damage response (DDR), cell proliferation, and cell cycle, as the most significantly downregulated biological processes upon depletion of *MALAT1* in LNCaP-abl cells (**Figure 3A,** and **Supplemental Table S2**). Likewise, an overlapping Metascape enrichment network of pathways also suggests that multiple biological processes associated with the DNA repair pathway and cell-cycle phase transition were the most significantly downregulated pathways in *MALAT1* depleted cells (**Supplemental Figure S2A).** Consistent with these findings, Gene Set Variation Analysis (GSVA) using three PCa cohorts (GSE35988 (29) GSE77930 (41), and GSE3325 (42)) exhibited significant enrichment (FDR<0.05) of the gene signatures associated with DDR and G_1_-S phase transition in metastatic PCa (**Supplemental Figure S2B**). Notably, a strong positive correlation (ρ⩾0.35) between *MALAT1* expression and DDR gene signature (retrieved from mSigDB) was also observed in these PCa datasets (**Figure 3B, and Supplemental Figure S2C).** To examine the clinical significance of the *MALAT1*-associated DDR genes, we performed Kaplan–Meier survival analysis for recurrence-free survival using the RNA-seq data of PCa patients from the TCGA-PRAD cohort categorized into two groups based on the median expression of *MALAT1* and the selected DDR genes. Intriguingly, patients with *MALAT1*-low and DDR-low signatures showed higher recurrence free survival probability compared with *MALAT1*-high and DDR-high patients (**Supplemental Figure S2D)**.

**Figure 3:**
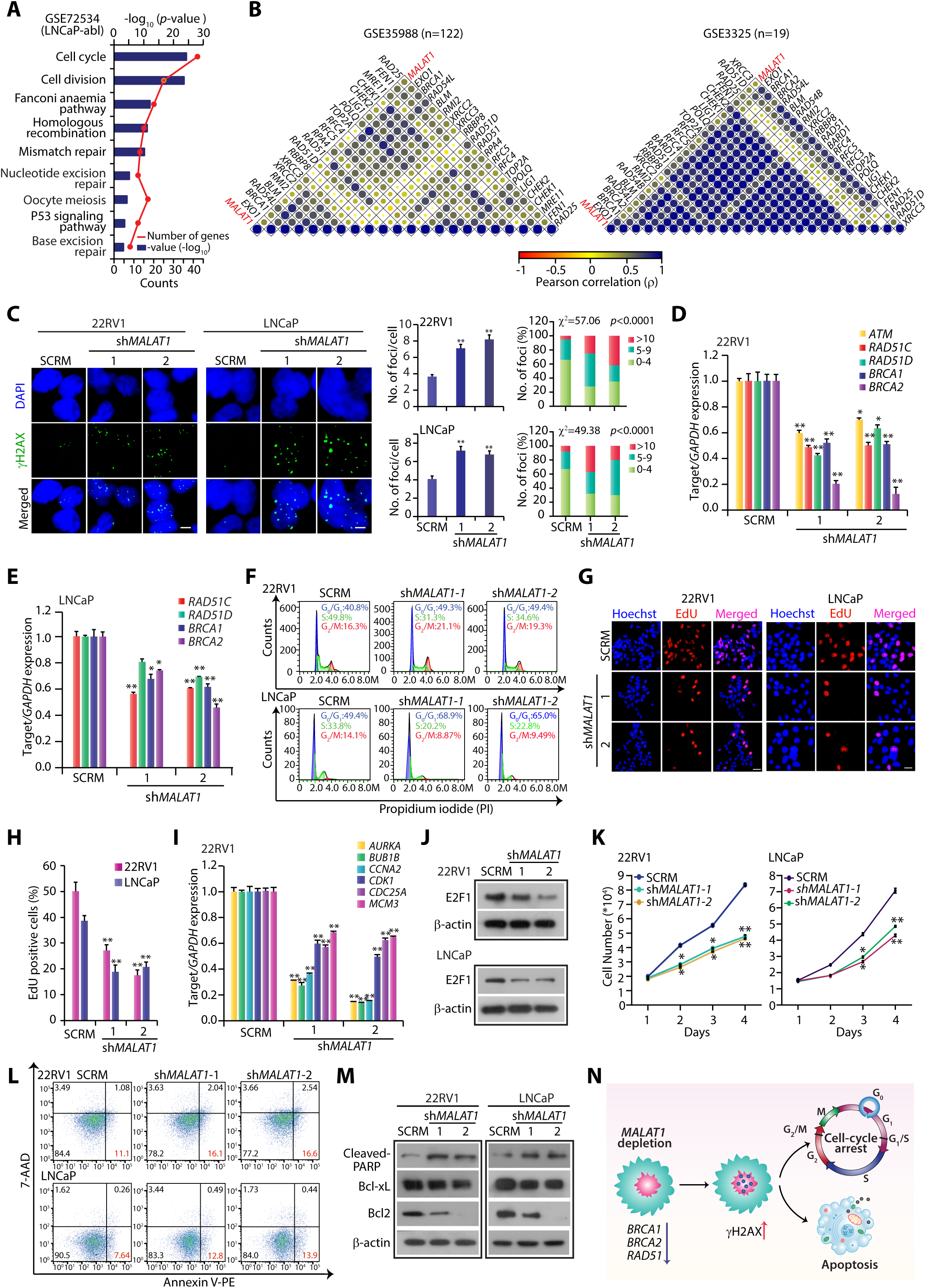
*MALAT1* depletion impairs homologous recombination-mediated DSB repair in prostate cancer. **A.** DAVID analysis shows biological pathways downregulated in LNCaP-abl-*siMALAT1* cells compared to LNCaP-abl-siCTL. Bars represent –log_10_ (*p-*values) and the frequency polygon (line in orange) represent the number of genes. **B.** Correlogram depicting Pearson correlation coefficient (ρ) between DNA repair associated genes and *MALAT1* in prostate cancer patient specimens (GSE35988 and GSE3325) (FDR adjusted, *p*<0.05). Correlation coefficients are expressed by shades of red and blue, and the size of dots is proportional to the strength of the correlation. Representative genes are marked on the sides of the correlogram. **C.** Representative confocal images for γH2AX foci (green) in SCRM control and *MALAT1* ablated PCa cells. The nucleus was visualized by DAPI (blue). The scale bar indicates 10 μm. Quantification of the number of γH2AX-positive foci (left panel) and bar plot showing the percentage of cells with the indicated number of foci/nuclei (right panel). The *p*-value for the Chi-Square test is indicated. **D.** Quantitative RT-PCR analysis depicting expression of the homologous recombination genes in 22RV1-sh*MALAT1* and SCRM control cells. Expression for each gene was normalized to *GAPDH*. **E.** Same as **D**, except LNCaP-sh*MALAT1* and SCRM control cells, were used. **F.** Flow cytometry analysis for accessing the cell cycle distribution by propidium iodide (PI) DNA staining assay in the same cells as in **C**. Percentage of cells in each phase was calculated using FlowJo software. **G.** Representative images depicting EdU incorporation in the same cells as in **C**. Nuclei were stained with Hoechst and the scale bar is 20 µm. **H.** Bar graph showing quantification of EdU uptake in the indicated cells. **I.** QRT-PCR analysis showing expression of genes associated with G1- and S-phase of the cell cycle in 22RV1-sh*MALAT1* and 22RV1-SCRM cells. The expression level for each gene was normalized to *GAPDH*. **J.** Immunoblot showing the change in the expression of E2F1 in the same cells as in **C**. β-actin was used as an internal control. **K.** Line graph showing cell proliferation assay using the same cells as in **C** at the indicated time points. **L.** Flow cytometry-based apoptosis assay using annexin V-PE and 7-ADD staining in the same cells as in **C**. Percentage of apoptotic cells was calculated using FlowJo software. **M.** Immunoblots showing expression of apoptosis markers in the same cells as in **C**. β-actin were used as an internal control. **N.** Schematics illustrate that *MALAT1* functions as a novel regulator of homologous recombination and plays crucial role in the maintenance of genome stability in prostate cancer. *MALAT1* facilitates the repair of double-strand breaks by modulating the expression of genes involved in DNA damage recognition and homologous recombination, thereby promoting cell cycle progression and accelerating cancer growth. Whereas *MALAT1* depletion leads to a decrease in the expression of several DDR genes and results in DSB accumulation which in turn induces cell-cycle arrest and instigates apoptosis. Experiments were performed with n= 3 biologically independent samples; data represents mean ± SEM. For panels, C, D, E, H, and I one-way ANOVA with Dunnett’s multiple-comparisons posthoc test was applied while the χ2 test was used for panel C. ∗*p*≤ 0.05 and ∗∗*p ≤* 0.001.

Subsequently, we examined the frequency of double-strand brakes (DSBs) in *MALAT1* silenced cells by examining the phosphorylation of H2AX on serine 139 residue (γH2AX). A marked increase in the abundance of γH2AX foci was observed in *MALAT1* deficient 22RV1 and LNCaP cells (**Figure 3C**), indicating that loss of *MALAT1* results in the accumulation of damaged lesions. Notably, RNA-seq data of *MALAT1* depleted LNCaP-abl cells exhibit a marked decrease in the expression of several genes that encode the proteins involved in the HR pathway (**Supplemental Figure S2E**). Likewise, 22RV1-sh*MALAT1* cells showed a robust decrease in the expression of key HR genes, such as *BRCA1, BRCA2, ATM, RAD51C,* and *RAD51D,* while a marginal decrease was observed in LNCaP-sh*MALAT1* cells (**Figure 3, D and E**); suggesting that *MALAT1* modulates expression of several HR genes and plays a crucial role in the maintenance of genome integrity.

On accumulation of DSBs, the cellular homeostatic mechanisms either obstruct the cell cycle to fetch additional time to repair the lesions or trigger apoptosis, if the damage is irreparable (43, 44). Besides, the expression of several HR proteins like RAD51 and BRCA1 is high during the S or G_2_ phase of the cell cycle, suggesting that a decrease in these HR proteins might influence the cell cycle progression (45, 46). We, therefore, examined the cell cycle distribution profile by performing propidium iodide (PI) staining. A robust increase in the G_1_ population with a concomitant decrease in the S phase cells was observed in *MALAT1* silenced 22RV1 (∼15%) and LNCaP cells (∼12%) (**Figure 3F)**. Likewise, the 5-ethynyl-2′-deoxyuridine (EdU) incorporation assay showed a robust decrease in the S-phase population upon *MALAT1* depletion in 22RV1 and LNCaP cells (**Figure 3, G and H)**. Remarkably, a significant decrease in the expression of genes associated with the cell cycle was observed in sh*MALAT1* PCa cells (**Figure 3I, and Supplemental Figure S3A**). Corroborating with this, a marked decrease in the expression of genes encoding for the proteins involved in G_1_-S and G_2_-M phase transition, such as *cyclin A2 (CCNA2), cyclin B (CCNB), cyclin-dependent kinases (CDKs), centromere proteins,* and *minichromosome maintenance (MCM2-8)* was also observed in the transcriptome profiles of *MALAT1* silenced LNCaP-abl cells compared to control (**Supplemental Figure S3B**). Similarly, a strong positive correlation (ρ⩾0.4) between *MALAT1* and cell-cycle associated genes was observed in PCa patient specimens (GSE35988 and GSE3325) (**Supplemental Figure S3C**), suggesting that *MALAT1* plays a pivotal role in cell cycle progression. We next examined the expression of E2F1, the major determinant of G_1_-S phase transition (47). Notably, a remarkable decrease in E2F1 levels was observed upon *MALAT1* depletion in PCa cells (**Figure 3J**). In agreement with this, *MALAT1* ablation markedly reduced cell proliferation in both 22RV1 and LNCaP cells **(Figure 3K).**

Since *MALAT1* depletion deregulates the cellular repair machinery, we speculated that the accumulation of DNA lesions might instigate apoptosis. So, next, we performed annexin-V and 7-AAD (7-amino-actinomycin D) staining and observed a robust increase in the number of late apoptotic cells in *MALAT1* depleted PCa cells (**Figure 3L)**. Likewise, levels of cleaved PARP, an early hallmark of apoptosis, also increased upon silencing *MALAT1* in both the cell lines (**Figure 3M**). Furthermore, a robust decrease in the anti-apoptotic Bcl-2 family proteins (BCL2 and Bcl-xL) was also observed (**Figure 3M**), establishing that the loss of *MALAT1* triggers apoptosis. Taken together, these findings firmly establish that *MALAT1* acts as a master regulator that modulates homologous recombination, cell cycle progression, and apoptosis in prostate cancer (**Figure 3N**).

### Attenuating *MALAT1* sensitizes prostate cancer cells to PARP inhibitors

Cancer cells with dysfunctional HR pathways rely heavily on PARP enzymes for removing the damaged lesion and ensuring their survival. Any additional pharmacological assault with PARP inhibitors or DNA damaging agents such as cisplatin, oxaliplatin, and carboplatin drives accumulation of DNA lesions and eventually induces cell death (48, 49). Since attenuation of *MALAT1* contrives HR deficiency in PCa, we speculated that the *MALAT1* deficient cells would be more vulnerable to chemotherapeutic agents that target DNA repair. To examine this, the *MALAT1* silenced 22RV1 and LNCaP cells and their respective SCRM control were treated with olaparib, an FDA-approved PARPi, and analyzed for any change in the oncogenic properties. Interestingly, *MALAT1* silenced 22RV1 as well as LNCaP cells exhibited decreased cell viability compared to the control, while the effect was more pronounced (∼80%) in olaparib treated cells (**Figure 4A)**. Similarly, colony forming ability of *MALAT1* silenced 22RV1 and LNCaP cells was significantly impaired in the presence of olaparib (**Figure 4, B and C**), indicating that *MALAT1* depletion augments olaparib activity. As a *MALAT1* depletion markedly enhances cellular sensitivity to PARPi, we next investigated the effect of olaparib treatment on cell-cycle profile in *MALAT1* silenced PCa cells by performing EdU staining. *MALAT1* knockdown cells upon olaparib treatment show a decrease in the number of S-phase population compared to the SCRM control in both the cell lines indicating dysregulation of the cell cycle (**Figure 4, D and E**).

**Figure 4:**
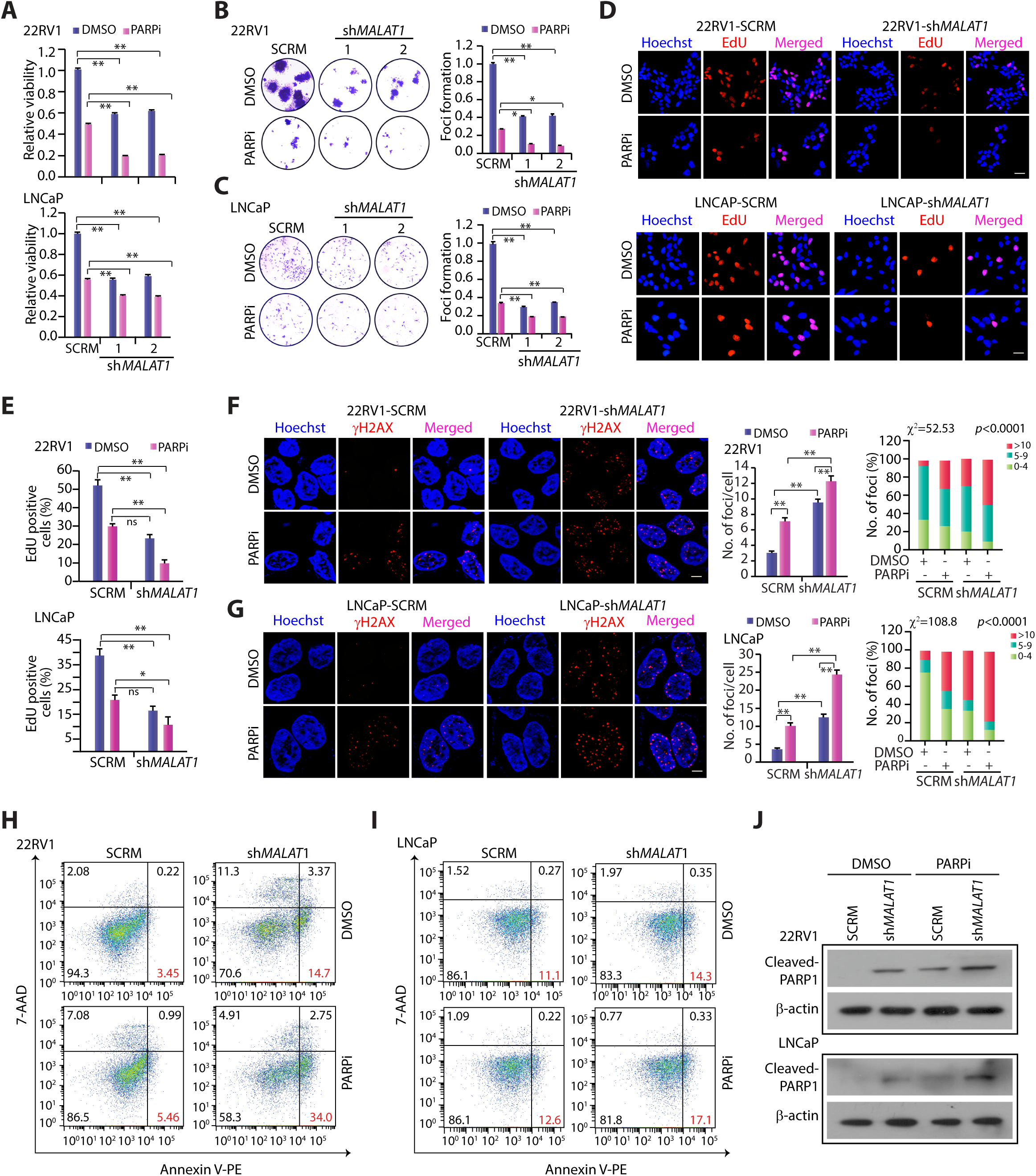
*MALAT1* depletion confers sensitivity to PARP inhibitors. **A.** Bar graph showing relative decrease in cell viability in olaparib treated (10μM) *MALAT1* silenced and SCRM control PCa cells. The drug was replenished every 24 hours for 2 days. **B.** Foci formation assay using 22RV1-sh*MALAT1* and SCRM control cells treated with olaparib (5μM) or vehicle control for 15 days. Inset showing representative images of foci. **C.** Foci formation assay using LNCaP sh*MALAT1* and SCRM control cells treated with olaparib (2μM) or vehicle control for 15 days. Inset showing representative images of foci. **D.** Representative confocal images for EdU uptake in the sh*MALAT1* and SCRM control PCa cells treated with olaparib (10μM) treatment for 48 hours. The scale bar indicates 50 μm. **E.** Bar graph showing quantification of EdU staining following 48-h treatment with olaparib in the indicated cells. **F.** Representative confocal images for γH2AX foci (red) in the same cells as in **B** upon olaparib (10μM) treatment for 48 hours. The nucleus was visualized by Hoechst 33342 (blue). The scale bar indicates 10 μm. Quantification of the number of γH2AX-positive foci (left panel) and bar plot showing the percentage of cells with the indicated number of foci/cell (right panel) in the same cells. The *p*-value for the Chi-Square test is indicated. **G.** Same as **F**, except LNCaP-SCRM and LNCaP-sh*MALAT1* cells. **H.** Flow cytometry-based apoptosis assay using annexin V-PE and 7-ADD staining in the same cells as in **B** upon olaparib (10μM) treatment for 48 hours. The percentage of the apoptotic cells population was calculated using FlowJo software. **I.** Same as **H**, except LNCaP-SCRM and LNCaP-sh*MALAT1* cells. **J.** Immunoblot showing a change in the expression of cleaved PARP in the same cells as in **A** upon olaparib (10μM) treatment for 48 hours. β-actin was used as an internal control. Experiments were performed with n = 3 biologically independent samples; data represents mean ± SEM. For panels A-E one-way ANOVA with Dunnett’s multiple-comparisons posthoc test was applied while the χ2 test was used for panels F and G. ∗*p* ≤ 0.05 and ∗∗*p*≤ 0.001.

Further, to assess that *MALAT1* depletion impairs the DNA repair system and sensitizes PCa to olaparib, we examined the frequency of γH2AX foci in *MALAT1* silenced cells following olaparib treatment. *MALAT1* depleted PCa cells exhibited almost two-fold higher γH2AX foci per cell in olaparib treated group compared to SCRM control (**Figure 4, F and G),** suggesting that *MALAT1* deficiency exacerbates olaparib induced DNA damage in PCa cells. Since the accumulation of DNA lesions instigate apoptosis, olaparib treated 22RV1-sh*MALAT1* and 22RV1-SCRM cells were further examined for cell death by AnnexinV-7AAD staining. A marginal increase in the late-apoptotic population was observed in *MALAT1* ablated 22RV1 and LNCaP cells compared to their respective SCRM controls, while upon olaparib treatment the number of apoptotic cells was markedly increased (**Figure 4, H and I**). Consistent with this, the level of cleaved PARP was increased in olaparib treated sh*MALAT1* - cells compared to SCRM control (**Figure 4J**). Collectively, our results provide compelling proof that *MALAT1* depletion enhances the sensitivity towards PARPi, hence targeting both key drivers will be a promising therapeutic approach for the treatment of advanced stage PCa patients, who often show resistance to conventional chemotherapies.

### *MALAT1* show positive association with neuroendocrine markers in castrate-resistant prostate cancer

Recently, it has been shown that NEPC patients exhibit higher expression of HR genes which play a crucial role in cell survival and cancer progression (50). Since higher expression of *MALAT1* has been reported in CHGA-positive NE tumors (51), we next sought to examine the functional significance of *MALAT1* in NE transdifferentiation. We analyzed publicly available RNA-seq data (GSE126078) consisting of transcriptome profiles of 98 metastatic tumors collected from 55 mCRPC patients (52) and discovered a positive association between *MALAT1* and NE genes. About 88% of NE tumors show higher levels of *MALAT1* (16 of 18), of which approximately 55% (10 of 18) also express HR genes (**Figure 5A**). Based on the expression pattern of NEPC markers, HR genes, AR, and its targets, we classified these NE tumors into two subtypes: a tumor cluster **AR^-^/ HR^+^/ NE^+^,** which manifests higher expression of *MALAT1*, HR genes, and NEPC genes, but lower expression of AR and PSA accounting ∼55% (10 of 18) of the NE-positive cases; and another cluster, namely **AR^+^/ HR**^-^/ **NE^+^** show co-expression of AR and its targets, and NEPC markers but lower expression of HR genes accounting 45% (8 of 18) of the cases.

**Figure 5:**
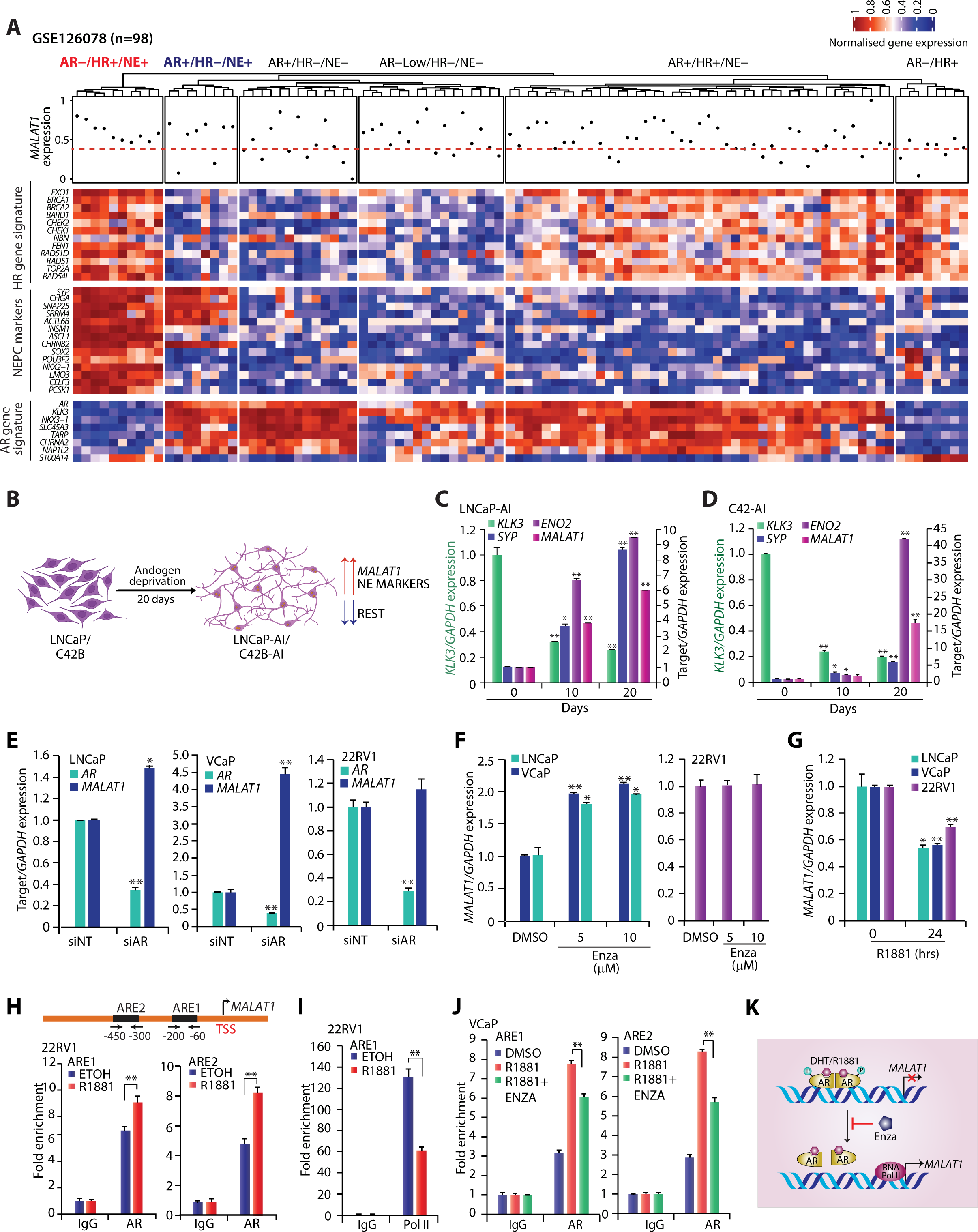
*MALAT1* positively associate with neuroendocrine prostate cancer. **A.** Heatmap depicting expression of HR genes, NE markers, AR, and its targets in mCRPC patients (n=98) retrieved from GSE126078. The normalized gene expression values are expressed by the shades of blue and red. Representative genes are marked on the left side of the heatmap. The topmost annotation in the heatmap indicates the *MALAT1* expression. **B.** The schema describes the process of neuroendocrine differentiation in LNCaP and C4-2 cells cultured in an androgen-free medium for 20 days. **C.** QPCR data showing relative expression of *MALAT1* and NE markers using androgen deprived LNCaP-AI cells. **D.** Same as C, except C4-2-AI cells. **E.** QPCR depicting expression of *MALAT1* in AR-silenced PCa cells. **F.** QPCR data showing relative expression of *MALAT1* in PCa cells treated with the indicated concentration of enzalutamide. **G.** Relative expression of *MALAT1* in PCa cells stimulated with R1881 (10nM) for 24 hours. **H.** Schema showing AR binding motif obtained from JASPAR database. The bottom panel shows the genomic location for the ARBs on the *MALAT1* promoter (top panel). ChIP-qPCR showing recruitment of AR on the putative *MALAT1* promoter upon R1881 (10 nM) stimulation in 22RV1 cells (bottom panel). **I.** Same as in **H**, except total RNA Pol-II marks. **J.** ChIP-qPCR depicting AR recruitment on the putative *MALAT1* promoter in R1881 (10nM) stimulated VCaP cells treated with or without Enzalutamide (10µM). Experiments were performed with n = 3 biologically independent samples; data represents mean ± SEM. For panels C, D, F, and G one-way ANOVA with Dunnett’s multiple-comparisons posthoc test was applied while a two-tailed unpaired Student’s *t*-test was used for panels E, H, I and J. ∗*p* ≤ 0.05 and ∗∗*p*≤ 0.001.

Next to examine the involvement of *MALAT1* in NE-transdifferentiation, we cultured LNCaP and C4-2 cells (LNCaP derivative osteotropic cell line) in androgen-deprived conditions for 20 days (**Figure 5B)** as androgen withdrawal is known to induce NE transdifferentiation (11, 53). A remarkable increase in *MALAT1* expression and NE markers namely, *SYP* and *ENO2* was observed with a concomitant decrease in *KLK3* levels (**Figure 5, C and D**).

As most NEPC tumors are characterized by low or absence of canonical AR signaling, we categorized TCGA-PRAD dataset into two groups based on AR levels and noticed that patients with higher AR expression showed lower levels of *MALAT1* compared to the patients with low AR (**Supplemental Figure S4A).** We next sought to examine the effect of AR signaling on *MALAT1* expression; for this, we silenced AR in LNCaP, VCaP, and 22RV1 cells using small interfering RNA (siRNA), and quantified *MALAT1* expression. Interestingly, *AR*-silenced LNCaP and VCaP cells exhibit an increase in the expression of *MALAT1*, while no change was observed in 22RV1 cells possibly due to abundance of *MALAT1* transcript (**Figure 5E).** Alternatively, small molecule inhibitor (enzalutamide) mediated pharmacological inhibition of AR signaling also results in a robust increase in *MALAT1* expression in both LNCaP and VCaP cells, while no change was observed in 22RV1 cells (**Figure 5F)**. In line with this, LNCaP cells (GSE152254) (54) treated with a lower concentration (1μM) of enzalutamide also show increased expression of *MALAT1* as well as AR repressed genes, namely *DDC* and *OPRK1* (**Supplemental Figure S4B).** Conversely, a marked decrease in *MALAT1* expression was noticed in VCaP cells stimulated with a sub-physiological concentration of synthetic androgen, R1881 (1 nM; GSE71797, (55) (**Supplemental Figure S4B).** Similarly, a significant decrease in the expression of *MALAT1* was noted in LNCaP, VCaP, and 22RV1 cells stimulated with R1881 (10nM) (**Figure 5G).** Taken together, these results imply that *MALAT1* being an androgen repressed gene is highly expressed in NEPC.

To further determine the impact of AR transcription factor on *MALAT1* expression, we scanned the *MALAT1* promoter for the presence of androgen response elements (AREs) using three publicly available transcription factor binding prediction software i.e., JASPAR, Promo-allgen, and MatInspector. Several putative AREs were identified within the ∼1 kb region upstream of the transcription start site (TSS) of *MALAT1* (**Figure 5H**). To confirm the binding of AR on the *MALAT1* promoter, we performed Chromatin immunoprecipitation (ChIP) using AR antibody in R1881-stimulated 22RV1 cells, interestingly a significant enrichment for AR-binding at two distinct AREs i.e., ARE1 and ARE2 were noted (**Figure 5H).** Moreover, reduced occupancy of RNA-polymerase II (RNA-Pol II) on the *MALAT1* promoter in R1881-stimulated 22RV1 cells indicates its transcriptional repression (**Figure 5I**). We next examined for any change in the recruitment of AR on the *MALAT1* promoter on restraining the androgen signaling by pharmacological inhibition using Enza. Significant enrichment of AR over input was observed on the *MALAT1* promoter in R1881 stimulated VCaP cells, while reduced occupancy was noted in Enza treated cells (**Figure 5J**), suggesting that AR negatively regulates *MALAT1* expression **(Figure 5K)**. Collectively our findings indicate that the *MALAT1* positively associates with the NE markers in prostate cancer.

### *MALAT1* promotes neuroendocrine transdifferentiation in prostate cancer

Having established an inverse association between AR signaling and *MALAT1*, we next sought to examine the functional implication of *MALAT1* in NE-transdifferentiation. For this, we first examined the expression of well-established NE markers and REST, the master transcriptional repressor of neuronal genes in *MALAT1* silenced 22RV1 cells. Intriguingly, 22RV1-sh*MALAT1* cells exhibit a remarkable increase in the REST levels (**Figure 6A**), with a concomitant decrease in the expression of key NE markers, namely, ENO2, SYP, and TUBB3, suggesting that *MALAT1* depletion hampers the expression of neuronal genes (**Figure 6A**). Further to confirm the involvement of *MALAT1* in the emergence of NE-like phenotype, we cultured LNCaP-SCRM and LNCaP-sh*MALAT1* cells in androgen-deprived condition for 30 days and performed phenotypic and molecular characterization of the cells. Intriguingly, *MALAT1* depletion effectively abrogated NE transdifferentiation as indicated by the absence of neuron-like projections in LNCaP-sh*MALAT1*-AI cells, while LNCaP-SCRM*-*AI attains neuronal projections (**Figure 6B**). Consistent with this, LNCaP-SCRM-AI cells upon androgen deprivation for 30 days also exhibited a decrease in REST and PSA levels with a concomitant increase in the expression of NE markers, EZH2, ENO2, and SYP (**Figure 6C).** Conversely, LNCaP-sh*MALAT1*-AI cells show a robust increase in the REST levels compared to LNCaP-SCRM-AI (**Figure 6C**). Moreover, androgen deprivation mediated increase in NE markers was more prominent in LNCaP-SCRM-AI compared to LNCaP-sh*MALAT1*-AI cells. Likewise, LNCaP cells transfected with other shRNA for *MALAT1* also exhibited a decrease in the expression of classical NEPC markers such as CHGA, EZH2, SYP, and ENO2 as compared to control cells (**Supplemental Figure S4C**), suggesting that *MALAT1* plays a pivotal role in promoting NE phenotype.

**Figure 6:**
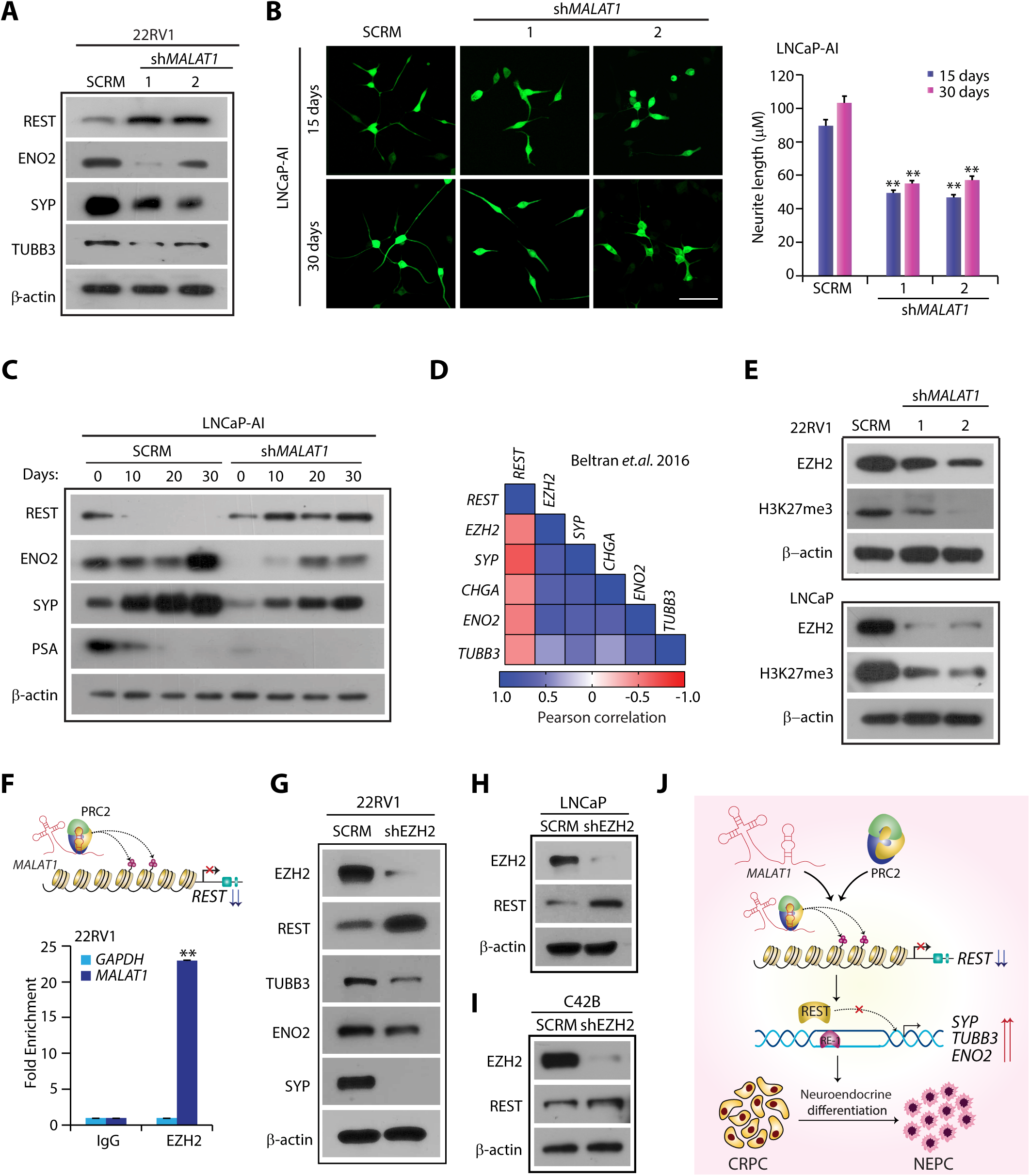
*MALAT1* plays a pivotal role in neuroendocrine trans differentiation. **A.** Immunoblot assay for REST, EZH2, TUBB3, ENO2, and SYP in 22RV1-sh*MALAT1* and SCRM control cells. β-actin was used as an internal control. **B.** Representative images for neurite outgrowth in androgen-deprived LNCaP-sh*MALAT1* cells and SCRM cells (left). Bar plot showing the length of neurite outgrowth of the indicated cells (right). The scale bar indicates 50 μm. **C.** Immunoblot analysis for NE markers and PSA using same cells as in **B**. **D.** Correlogram depicting Pearson correlation coefficient (ρ) between NE genes and *REST* in aggressive prostate cancer patient samples from Beltran cohort, 2016 (FDR adjusted, *p*<0.05). Correlation coefficients are expressed by shades of red and blue and representative genes are marked on the sides of the correlogram. **E.** Immunoblot depicting expression of EZH2 and H3K27 trimethylation levels in sh*MALAT1* and SCRM PCa cells. β-actin was used as an internal control. **F.** RIP assay followed by qPCR showing associations of *MALAT1* with EZH2 as compared to IgG (control antibody) in 22RV1 cells. **G.** Immunoblot depicting expression of REST, EZH2, and NE markers in 22RV1-SCRM and 22RV1-sh*EZH2* cells. β-actin was used as an internal control. **H.** Immunoblot depicting expression of REST and EZH2 in LNCaP-SCRM and LNCaP-sh*EZH2* cells. β-actin was used as an internal control. **I.** Same as H, except C4-2B cells, were used. **J.** Schematic depicting that role of *MALAT1* in inducing neuroendocrine trans-differentiation by epigenetically silencing of REST via epigenetic modulator EZH2 in PCa cells. Experiments were performed with n = 3 biologically independent samples; data represents mean ± SEM. For panel B one-way ANOVA with Dunnett’s multiple comparisons, a posthoc test was applied. ∗*p* ≤ 0.05 and ∗∗*p*≤0.001.

Since *MALAT1* depletion leads to a robust increase in the REST levels, we speculated that the elevated REST disrupts NE transdifferentiation in *MALAT1* silenced cells. Thus, we generated stable 22RV1 cells overexpressing REST and examined the expression of NE markers. Interestingly, we noticed a robust decrease in the expression of SYP and CHGA (**Supplemental Figure S4D**). Likewise, a significant decrease in SYP and TUBB3 expression was noticed, when we restored REST levels in 22RV1 cells using a pharmacological inhibitor for CK1 inhibitor (iCK1, D4476, 56,57) (**Supplemental Figure S4E**). Conversely, shRNA-mediated knockdown of REST in LNCaP and C4-2B cells led to a robust increase in SYP levels (**Supplemental Figure S4F**). Further, to examine the association between REST and archetypal NEPC markers, we calculated Pearson correlation in advanced stage PCa patients using the Beltran cohort, 2016 (17) and found a negative correlation between REST and *SYP, CHGA, ENO2,* and *TUBB3* (**Figure 6D)**, suggesting that elevated levels of REST in *MALAT1* silenced cells might perturb the expression of neuronal genes.

To identify a plausible regulator of REST in NEPC, we next examined the association of REST with previously identified transcription factors that are essential for NE transdifferentiation. Interestingly, we found that EZH2, a member of polycomb repressor complex 2 negatively associates with REST in NEPC patients (**Figure 6D)**. Moreover, EZH2 is known to physically interact with several lncRNAs which acts as a molecular scaffold for the PRC2 and facilitates its recruitment to specific genomic loci to modulate the expression of the target gene by enhancing tri-methylation marks on the lysine 27 residue of histone H3 (H3K27me3) (58, 59). Furthermore, *MALAT1* is known to recruit EZH2 to the promoter of epithelial genes such as CDH1 and PCDH10 (60, 61). Therefore, we speculated that *MALAT1* interacts with EZH2 to modulate the expression of REST in advanced-stage PCa. To ascertain the oncogenic collaboration between *MALAT1* and EZH2, we first examined their correlation in PCa patient samples using two publicly available datasets, namely GSE35988 (29) and GSE77930 (41). Interestingly, *MALAT1* and *EZH2* show a positive association (ρ*⩾*0.05; **Supplemental Figure S4G**) in all three datasets. In line with this, a marked decrease in EZH2 expression, as well as H3K27me3 marks was noted in *MALAT1* silenced 22RV1 and LNCaP cells (**Figure 6E**). Further to confirm the association between *MALAT1* and EZH2, we performed RNA immunoprecipitation (RIP) assay using anti-EZH2 antibody, intriguingly a remarkable enrichment in *MALAT1* transcript (∼22 folds) was observed with EZH2 antibody compared to IgG control (**Figure 6F**), confirming direct interaction between *MALAT1* and EZH2. To further confirm that the epigenetic regulator EZH2 modulates the expression of REST in advanced stage PCa, we established stable EZH2-silenced 22RV1, LNCaP, and C42B cells using shRNA. A remarkable increase in the REST levels with a concomitant decrease in the expression of NE markers was observed in EZH2 depleted cells (**Figure 6, G-I**). Collectively, our findings show that tumors with higher expression of *MALAT1* have a greater tendency to acquire NE phenotype by PRC2 complex mediated downregulation of REST (**Figure 6J**).

## DISCUSSION

In this study, we demonstrate that lncRNA *MALAT1* plays a vital role in maintaining genome integrity and helps escaping PCa cells from chemotherapeutic drugs by enhancing the expression of HR genes. Furthermore, meta-analysis of gene expression data from three independent PCa datasets showed a strong positive correlation between *MALAT1* and core HR genes. In addition, HR genes, as well as *MALAT1*, were noted to be enriched in metastatic PCa patients with higher Gleason grade, suggesting that oncogenic coordination might contribute to aggressive disease and therapy failure. Moreover, *MALAT1* knockdown results in the accumulation of DSB lesions with a concomitant depletion in several HR proteins, including *RAD51, BRCA1* and *BRCA2*. In consonance with our findings, recent reports demonstrate that *MALAT1* depletion enhances sensitivity toward chemotherapeutic drugs like docetaxel (62), oxaliplatin (63), cisplatin (64), temozolomide (65), cytarabine (66), and 5-fluorouracil (67). While Hu et. al. showed that *MALAT1* plays a critical role in regulating the alternative NHEJ pathway by interacting with PARP1 and LIG3 (68). These findings suggest that *MALAT1* is an important molecular modulator of the HR pathway that may augment chemosensitization.

Furthermore, we show that *MALAT1* modulates the expression of several essential genes needed for G_1_/S transition, including *CDK1*, *cyclin A2,* and *Cdc25A*, and its depletion results in G_1_/S transition arrest with a concomitant decrease in the S-phase population. In agreement with this, Vidisha et. al. also demonstrated that the *MALAT1* levels vary through different phases of the cell cycle wherein G1/S and M phase exhibits its higher expression. They showed that *MALAT1* depletion perturbs the cell cycle machinery by suppressing the expression of genes involved in G_1_/S and mitotic progression, thus supporting our findings that *MALAT1* plays a crucial role in cell cycle regulation (69).

Our findings suggest that the HR deficiency induced by *MALAT1* depletion phenocopies “BRCAness”, and exhibits contextual synthetic lethality with clinically approved DNA repair inhibitors such as Olaparib. Of note, *MALAT1* depletion exhibits anti-proliferative effect and delays the resolution of γ-H2AX foci in HR deficient PCa cells upon olaparib treatment. Overall, our findings suggest that *MALAT1* ablation may provide a considerable therapeutic benefit to advanced stage PCa patients when combined with PARPi or any other DNA damaging agents. Nevertheless, many critical questions remain unresolved, including the molecular mechanism by which *MALAT1* regulates the expression of HR genes. Our analysis of the publicly available CHART-seq data (GSE58444) (70) revealed that *MALAT1* was not enriched on the genomic locus of key HR genes, namely *BRCA1, BRCA2,* and *RAD51* (data not shown), suggesting that *MALAT1* influences HR gene expression indirectly probably through a transcription factor or miRNA, which warrants an in-depth investigation. Despite these limitations, this is the first study to demonstrate the direct molecular interplay between a lncRNA and DDR pathway in PCa.

It has been shown that CHGA-positive NE tumors exhibit higher expression of *MALAT1* (51), which is in line with our findings wherein *MALAT1* is involved in lineage plasticity and facilitates NE transdifferentiation. This data was further strengthened by a recent report which demonstrates that *MALAT1* expedites neurite outgrowth in neuroblastoma-derived Neuro-2a cells while its knockdown results in neurite outgrowth defects (71). Yet another study showed that *MALAT1* modulates synapse formation by regulating the expression of key neuronal genes such as *neuroligin1 (NLGN1*) and *SynCAM1* (*72*). Collectively, these independent findings reaffirm the functional implication of *MALAT1* in NE transdifferentiation.

Of note, our research deciphered the unknown molecular connections between *MALAT1* and epigenetic modulators, namely, REST and EZH2. Although their functional role in NE development is widely recognized, their molecular association has not been established yet. Our findings reveal a negative association between REST and EZH2 in NEPC patients while, *MALAT1* is known to facilitate the recruitment of EZH2 on its target genes, which in turn establishes H3K27me3 repressive marks and suppresses the expression of its target genes (58, 59). Thus, given the central role played by EZH2 in NEPC, we conjecture that *MALAT1* acts as a transacting factor that suppresses REST expression by interacting with the EZH2, leading to enhanced expression of NE markers.

In consonance with our findings, Zhang et. al. also reported higher levels of HR genes in NEPC patients compared to CRPC cases and showed that suppression of key HR genes such as *BRCA1* and *RMI2* led to reduced cell proliferation and an increase in sub G_1_ population (50). Nevertheless, another study showed that PARPi impedes NE transdifferentiation and suppresses tumor growth in therapy-induced NEPC (73). These studies along with ours provide a molecular rationale for targeting *MALAT1* in combination with PARPi for NEPC patients, for whom effective treatment options are not available. Therefore, it is important to further investigate how *MALAT1* modulates the dynamic process of NE transdifferentiation.

Collectively, our findings suggest that *MALAT1* escapes tumor cells from anticancer agents by initiating DNA repair pathways and point toward a possible therapeutic vulnerability that can be exploited by targeting *MALAT1* along with PARP inhibitors. We also discovered previously unknown molecular connections between *MALAT1* and neuroendocrine transdifferentiation, which could be targeted as a potential therapeutic avenue for patients with this lethal subtype. Conclusively, our findings provide a compelling rationale for conducting clinical trials in patients with advanced-stage disease and investigate the safety and efficacy of a combination therapy consisting of *MALAT1* antisense oligonucleotides/GAPmers or small molecular inhibitors with DNA damaging agents like PARPi or cisplatin.

## METHODS

### In silico data processing and computational analysis

#### Microarray analysis

The gene expression datasets, namely GSE6919, GSE6752, and GSE35988 were downloaded from the GEO database (https://www.ncbi.nlm.nih.gov/geo/), each of which contains expression profiles for localized PCa and CRPC samples. The differentially expressed genes (DEGs) in CRPC patients were identified using the "limma" package in R with the cutoff criterion of adjusted *p*-value 0.05 and log_2_ fold change |FC| > 0.6 (74). The commonly elevated genes in CRPC samples in the aforementioned cohorts were identified by Venn analysis using the webtool (https://www.ncbi.nlm.nih.gov/geo/). Samples were sorted based on tissue type and *MALAT1* expression was plotted (log_2_ (normalized count)) using GraphPad prism version 7.0.

#### Gene co-expression analysis

The pairwise Pearson’s correlation coefficient (ρ) between *MALAT1* and DDR/cell-cycle genes in PCa cohorts namely GSE35988, GSE3325, and GSE77930 was examined using the “corrplot” package in R. *P*-value < 0.05 was considered statistically significant threshold. Further, gene set variation analysis (GSVA) (75) was performed to investigate the variations in the activation status of the DDR pathway and G_1_/S phase transition in different clinical stages of PCa. The gene sets for "DDR pathway" and "G_1_/S phase transition" were downloaded from the MSigDB database (https://www.gsea-msigdb.org/gsea/msigdb/), and statistical significance was defined as an adjusted *p* ≤ 0.05. Ultimately, an R package named “pheatmap” was used to demonstrate the enrichment of the aforementioned gene sets in each group and the gene expression of *MALAT1* (76).

For the publicly available datasets where VCaP (GSE71797) and LNCaP cells (GSE152254) were treated with R1881 and enzalutamide (Enza) respectively, heatmaps (77) were generated using gplots’s heatmap.2 function.

#### Integrative analyses for TCGA-PRAD data

The HiSeq mRNA data for *MALAT1* and clinical information of TCGA-PRAD dataset were obtained from the UCSC Xena browser (https://xenabrowser.net). The gene expression data were log_2_ transformed. We analyzed the association of *MALAT1* with the routine clinicopathological parameters such as Gleason score, node status, and response to therapy. Further, the samples were stratified into two groups based on the median expression value of *MALAT1* wherein patients with expression values higher than the median were placed in the ‘*MALAT1* high*’* group, while patients with expression values lower than the median were placed in the ‘*MALAT1* low’ group. Further, Kaplan-Meier survival analysis was performed using GraphPad Prism 7 to calculate the 5-year survival probability and the log-rank test to detect significance.

#### RNA sequencing analysis

GEO accession numbers GSE72534 and GSE126078 were used to download RNA sequencing data in FASTQ format. FastQC sequence quality checks were performed on the raw reads before mapping it to hg38 human using TopHat v2.1.0. The Genomic Alignments Bioconductor software in R was used to calculate gene-level abundances (78). The edgeR Bioconductor program was used to determine the differential expression of transcript abundances (79). In order to identify differentially expressed genes, Benjamini and Hochberg’s procedure was used to calculate FDR-corrected p-values (FDR ⩽* 0.05).

The differentially downregulated genes in LNCaP-abl-sh*MALAT1* cells (log_2_ fold change ≤ −0.6) were then subjected to the DAVID bioinformatics platform (Database for Annotation, Visualization, and Integrated Discovery) to identify deregulated biological processes (*P*<0.05) (40). Further, functional enrichment analysis of biological pathways was generated using Metascape (http://metascape.org)). Term significance was assessed based on a *p*-value⩽0.01, minimum count of 3, and enrichment factor of >1.5. Within a cluster the most statistically significant term was chosen as its representative. To further determine the relationship among enriched terms, a network plot was generated by selecting a subset and connecting terms with a similarity of >0.3 by edges.

### Experimental methods

#### Cell lines culture conditions and authentication

Prostate cancer cell lines (22RV1, VCaP, LNCaP, PC3) and benign prostate epithelial cells (PNT2) were sourced from the American Type Culture Collection (ATCC). DU145, C42, and C42-B cells were generously gifted by Dr. Mohammad Asim, Department of Clinical and Experimental Medicine, Faculty of Health and Medical Sciences, University of Surrey, Guildford, UK. The cells were cultured according to ATCC recommended guidelines, in 37°C incubators with 5% CO2. For experiments performed in androgen-depleted conditions, RPMI without phenol red (Gibco) supplemented with Charcoal Stripped FBS was used. Cell lines were periodically tested for *Mycoplasma* contamination using the PlasmoTest mycoplasma detection kit (InvivoGen). The cell lines used in this study were authenticated by short tandem repeat (STR) profiling at the Life code Technologies Private Limited, Bangalore, and DNA Forensics Laboratory, New Delhi.

#### Lentiviral packaging

ViraPower Lentiviral Packaging Mix (Invitrogen) was used to generate viral particles for shSCRM/sh*MALAT1* constructs as previously described (80). Briefly, the shRNA constructs and packaging mix plasmids were transfected into HEK293FT cells and incubated for 60-72 h. The viral particles were then harvested and stored at −80 °C. To establish stable lines, the 22RV1 and LNCaP cells were infected with the collected lentiviral particles along with polybrene (hexadimethrine bromide; 8 µg/ml) (Sigma-Aldrich). Culture media was changed the next day and puromycin (Sigma-Aldrich) selection was started three days after infection.

#### Immunoblotting

The cells were lysed in radioimmunoprecipitation assay buffer (RIPA buffer) supplemented with phosphatase inhibitor cocktail set-II (Calbiochem) and protease inhibitor (VWR). The BCA assay kit was used to quantify the isolated protein samples (GE Healthcare). The proteins were then heat-denatured and resolved on SDS-PAGE. The proteins were then transferred to a polyvinylidene fluoride (PVDF) membrane (GE Healthcare), and the membrane was blocked with 5% non-fat dry milk for 1 hour at room temperature. The membrane was incubated with primary antibodies overnight at 4 °C with 1:1000 dilution-N-Cadherin (Abcam, ab98952), E2F1 rabbit (1:1000, 3742S), Cleaved PARP (CST, 9541), Bcl-xL (CST, 2764), BCl2 (CST, 762870), TUBB3 (CST, 5568), EZH2 (Abcam, ab191250), H3K27me3 (CST, 9733), 1:2000 diluted REST (Abcam, ab75785), PSA (CST, 5877), SYP (Abcam, ab32127), ENO2, 1:5000 diluted E-cadherin (CST, 3195) and β-actin (Abcam, ab6276). Following this, the membrane was washed thrice with 0.1% TBS-T and incubated for 2 hours at room temperature with horseradish peroxidase (HRP) conjugated secondary anti-mouse (Jackson ImmunoResearch Laboratories, Cat # 115-035-003) or anti-rabbit antibody (Cat # 111-035-144). Membranes were washed again with 0.1% TBS-T buffer, and the signals were detected through an enhanced chemiluminescence system (Thermofisher) as directed by the manufacturer.

#### Cell migration assay

Transwell Boyden chamber with an 8μm pore size (Corning) was employed for migration assay. The upper compartment was seeded with 1*10^5^ cells suspended in serum-free culture medium, while the lower compartment was filled with conditioned media supplemented with 20% FBS. After a 24-hour incubation at 37°C, the cells were fixed in paraformaldehyde (4%) in 1X PBS and stained with crystal violet (0.5%) w/v). Cells attached to a Transwell filter were de-stained with 10% v/v acetic acid. An upright Nikon microscope was used to examine and photograph migrating cells.

#### Real-time quantitative PCR

Total RNA from cells was extracted using TRIzol (Ambion) and reverse transcribed to cDNA using the First Strand cDNA synthesis kit (Genetix Biotech Asia Pvt. Ltd) according to the manufacturer’s instructions. The expression of target RNA wasvalidated by performing qPCR in triplicates using SYBR Green PCR Master-Mix (Applied Biosystems), primers mentioned in **Supplemental Table S3**. *GAPDH* was used as an internal reference and relative expression of the target gene was calculated using the -ΔΔCt method.

#### Flow cytometry

Cells were grown to 80% confluence and dissociated using StemPro™ Accutase™ (ThermoFisher). The cells were washed twice and resuspended in phosphate-buffered saline with 5% FBS at 10^6^ cells/ml. Further, 100 μl of cell suspension was incubated with CD117-APC (c-KIT) antibody (Miltenyi Biotec, 130-098-207, 1:50), CD133-PE (PROM1) antibody (Miltenyi Biotec, 130-113-670, 1:50), and CD338-PE (ABCG2) antibody (Miltenyi Biotec, 130-105-010, 1:50) for 1 h on ice, then washed three times in phosphate-buffered saline with 5% FBS. The events were acquired on BD Beckman Coulter’s CytoFLEX platform and analyzed with FlowJo version 10.7.

#### Cell cycle distribution. MALAT1

silenced cells were fixed in 70% ethanol and stained with propidium iodide (PI) (BioLegend, Cat # 421301) as directed by the manufacturer’s protocol. The acquired events were analyzed using FlowJo software version 10.7’s built-in univariate model was used to assess cell cycle distribution.

#### Apoptotic assay. MALAT1

silenced cells were stained with the PE Annexin V Apoptosis Detection Kit I (BD Biosciences, Cat # 559763) following the manufacturer’s protocol. Quadrants were gated on GFP, Annexin (PE), and 7AAD (PerCP) channel dot plots using unstained, GFP, Annexin V (PE), and 7AAD (PerCP) single stained cells as controls. The cells were divided into four quadrants: lower left quadrant Annexin^−^/7AAD^−^ (viable), lower right quadrant Annexin^+^/7AAD^−^ (early apoptotic), and higher left quadrant Annexin^−^/7AAD^+^ (necrotic), and upper right quadrant Annexin^+^/7AAD^+^ (late apoptotic) cells were characterized. Further, flowJo software v10.7 was employed to analyze the data acquired on the BD FACS Melody Cell Sorter for each condition.

#### Prostatosphere assay

22RV1 shSCRM and 22RV1 sh*MALAT1* cells (1 × 10^4^) were plated in a serum-free stem cell medium consisting of DMEM-F12 (1:1, Invitrogen), 1X B27 (Invitrogen), 20 ng/ml EGF (Invitrogen), and 20 ng/ml FGF (Invitrogen). Every 3^rd^ day, the prostatospheres formed were dissociated into single-cell suspension, and then re-plated in fresh medium for multiple generations. The experiment was terminated after 12 days and bright-field images of the spheres were captured using Axio Observer Z1 inverted fluorescence microscope (Carl Zeiss) equipped with an Apotome device. Further, the prostatospheres formed in all the groups were counted and analyzed for sphere-forming efficiency using ImageJ software. The spheroids were also subjected to RNA isolation, as previously described (11, 81).

#### Cell proliferation assay

The cell proliferation was performed by plating 1*10^4^ cells per well in a 12-well plate and counting the cells at the indicated time points. Cells were treated with 10μM of olaparib against DMSO control and incubated for the indicated time points. Cells were counted at the indicated time points using a hemocytometer.

*Foci formation assay*. *MALAT1* depleted 22RV1 and LNCaP (2*10^3^) cells were cultured in serum-deprived conditions in RPMI-1640 medium (Gibco). Media was replenished after every 48 h with Olaparib (5µM for 22RV1 and 2μM for LNCaP) along with DMSO control. After 2 weeks the foci formed in each condition were fixed with 4% paraformaldehyde (PFA) and stained with crystal violet (0.1% w/v). While destaining was done using 10% glacial acetic acid and the absorbance was measured at 550 nm.

#### Immunofluorescence staining

Cells were seeded on sterilized glass coverslips in 4-well plates and incubated for 36 h. The cells were washed twice with PBS and then fixed with 4% paraformaldehyde (PFA) in PBS. The cells were washed with 1×PBS thrice and permeabilized with 0.3% Triton X-100 in PBS. Blocking was done using 5% normal goat serum (NGS) in PBS-T for 2 h at room temperature. The cells were then incubated with primary antibodies: c-Kit or CD117 (1:400, CST, 3308), CD44 (1:400, CST,3570), and γ-H2AX (1:100, CST, 2577) diluted in PBS-T, at 4 °C overnight. Cells were washed thrice times with PBS-T before being stained with DAPI (SigmaAldrich). The coverslips were then adhered to glass slides using an anti-fade Vectashield mounting medium (Vector Laboratories). Images were captured with a Carl Zeiss Axio Observer Z1 inverted fluorescence microscope fitted with an Apotome device. For neurite outgrowth measurement, 10 random areas of fields of LNCaP-SCRM and LNCaP-shMALAT1 cells grown in androgen-free conditions were imaged. The Simple Neurite Tracer (http://imagej. net/Simple Neurite Tracer) was used to measure neurite lengths.

#### Chemosensitivity assay

For determining the IC50 of drugs, 22RV1-shSCRM and -sh*MALAT1* cells (2 × 10^3^) were seeded in 96-well plates and treated with varying concentrations of doxorubicin and 5-fluorouracil for 48 h. Resazurin (Cayman Chemicals) was added to each well according to the manufacturer’s instructions and after 3 hours’ fluorescence was measured with emission-excitation at 590-530nm. The IC50 of the drugs was calculated by linear approximation regression of the percentage survival versus the drug concentration.

#### EDu uptake

*MALAT1* silenced PCa cells were stained with Click-iT EdU Cell Proliferation Kit for Imaging (Thermofisher, C10338) as directed by the manufacturer’s protocol. In brief, 1*10^5^ cells were seeded on glass coverslips in 12-well plates and incubated for 36 hours before being treated with 10 μM EdU for 2 hours at 37 ^O^C. Following that, the cells were fixed with 4% paraformaldehyde, permeabilized with 0.3 percent Triton X-100 in 1 PBS (PBS-T) for 10 minutes, then stained with EdU Azide Alexa Fluor 555, followed by Hoechst 33342 (Thermofisher, 62249). Vectashield mounting medium was used to adhere the coverslips to the glass slides (Vector laboratories). Images were taken with a Carl Zeiss Axio Observer Z1 inverted fluorescence microscope fitted with an Apotome device. ImageJ software was used for post-processing and quantification of the acquired images.

#### Chromatin immunoprecipitation (ChIP) assay

ChIP was performed as described previously (11, 80). Briefly, the cells were crosslinked with 1% formaldehyde for 10 min and then quenched with 125 mM glycine. The cells were lysed first in lysis buffer [1% SDS, 50 mM Tris-Cl (pH 8.0), 10 mM EDTA, protease inhibitor cocktail (Genetix) and phosphatase inhibitor cocktail (Calbiochem)], and then sonicated using Bioruptor (Diagenode) to get ∼500 bp DNA fragments. The sheared chromatin was incubated with 4 µg of anti-AR (CST, 5153) or isotype control antibody, rabbit IgG (Invitrogen, 10500C) at 4 °C. Simultaneously, Dynabeads coated with Protein G (Invitrogen) were blocked sheared salmon sperm DNA (Sigma-Aldrich) and bovine serum albumin (BSA) (HiMedia) overnight at 4 °C. Blocked beads were then incubated with the antibody-bound lysate to create antibody-bead conjugates. After washing the antibody-conjugated beads were eluted using elution buffer [1% SDS, 100mM NaHCO3, Proteinase K (Sigma-Aldrich), and RNase A (500µg/ml each) (Sigma-Aldrich)]. DNA was extracted using the phenol-chloroform-isoamyl alcohol method.

#### RNA Immunoprecipitation (RIP)

22RV1 cells were plated at ∼90% confluency 36 hours before harvesting cells. Briefly, the cells were washed twice with ice-cold 1X PBS and scraped in 1X PBS with a protease inhibitor. After this the cells were incubated in RIP buffer [50 mM Tris-HCl (pH 7.9), 0.25 M NaCl, 1% Nonidet P-40 (NP-40), 10 mM EDTA, protease inhibitor cocktail and RNase inhibitor] for 30 min. Further, cell lysate was obtained by centrifugation/n at 12,000 rpm for 10 min at 4°C and was incubated overnight with 4µg of EZH2 antibody or IgG. Simultaneously the Dynabeads (Protein G coated, Invitrogen) were pre-absorbed with 100µg/ml BSA at 4°C. The pre-absorbed beads were washed thrice with NT2 buffer [50 mM Tris-HCl (pH 7.4), 300 mM NaCl, 1 mM MgCl2, 0.05% Nonidet P-40 (NP40), 1 × PIC, RNase inhibitor] and then incubated with RNA-antibody adducts to form RNA-antibody-bead precipitates. Further, the beads were washed thrice with NT2 buffer, followed by DNase I digestion for 15 min at 37°C, and washed twice with NT2 buffer. Co-purified RNA was extracted by Trizol and cDNA was synthesized using the SuperScript kit from Puregene.

#### Statistics

For statistical analysis, unpaired two-tailed Student’s *t*-test, one-way ANOVA, or two-way ANOVA were employed, unless otherwise stated in the respective figure legend. *P*-value ⩽0.05 was considered significant wherein \**p*⩽0.05, \*\**p*⩽0.001 were used to denote significance. The error bars indicate the standard error of the mean (SEM) determined from at least three independent experiments.

## ACKNOWLEDGEMENTS

The authors thank the Indian Institute of Technology Kanpur for the infrastructure support. We are thankful to Prof. Jonaki Sen and Prof. Pradip Sinha for extending the use of their microscope facility. We thank all members of our laboratory for insightful discussions during the research, and Mahendra Palecha for technical assistance.

## AUTHOR CONTRIBUTIONS

A.Y. and B.A. conceptualized the study and designed the experiments. A.Y. and T.B. performed *in-vitro* studies. A.Y. and A.P. performed the gene expression studies and bioinformatics analysis. P.G. analyzed the CHART-seq data. A.V. and D.D. assisted in *in-vivo* experiments. A.Y. performed the statistical analysis. A.Y. and B.A. interpreted the data and wrote the manuscript. B.A. directed the overall project.

