## Supplement Figures for "Targeting *MALAT1* Augments Sensitivity to PARP Inhibition by Impairing Homologous Recombination in Prostate Cancer"

### Supplemental Figure 1

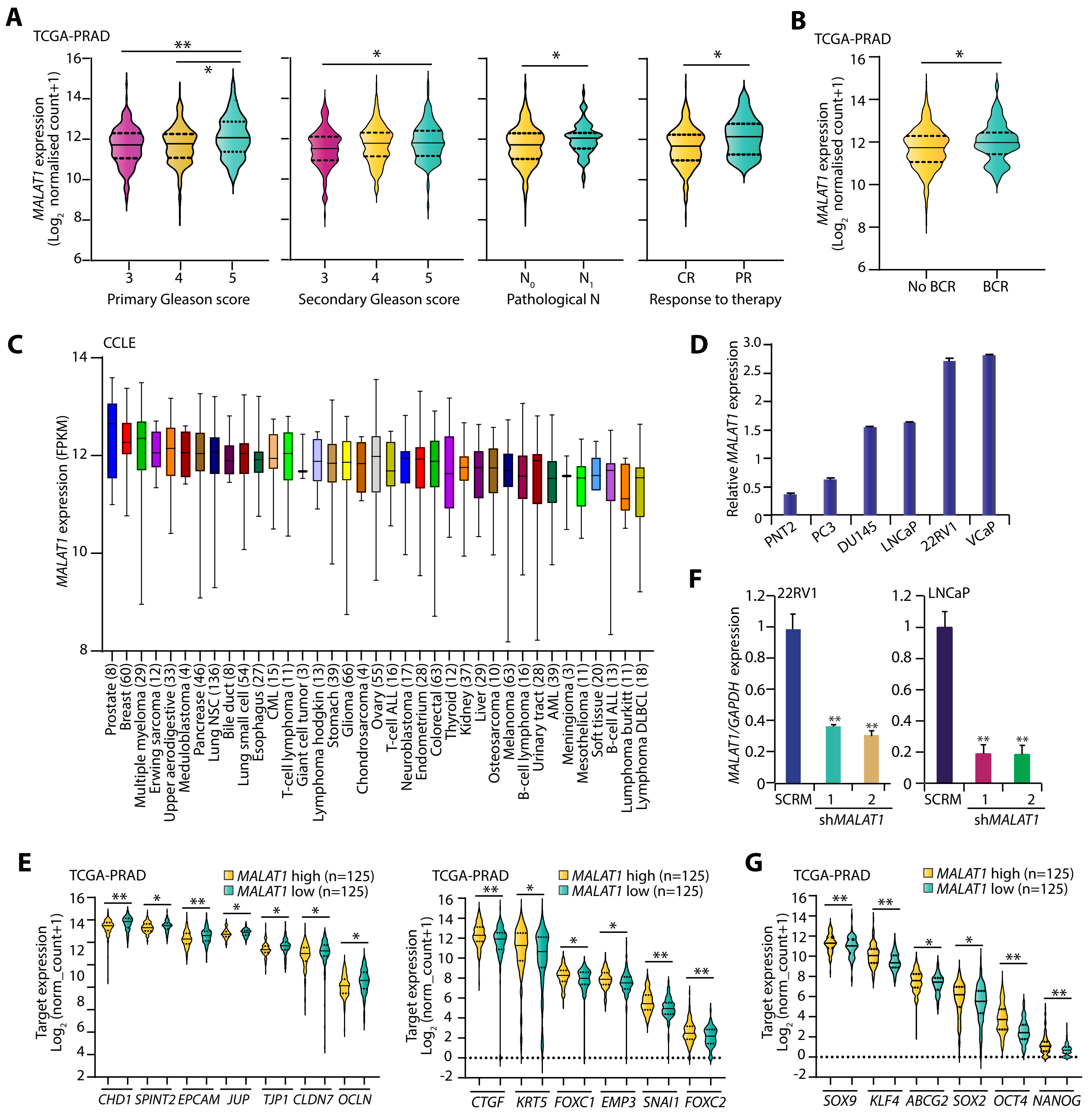


**Supplemental Figure 1: *MALAT1* is upregulated in prostate cancer and positively associates with aggressive clinical phenotype.**

**A.** Violin plot comparing the *MALAT1* expression in prostate cancer patients (n=499) categorized by varying primary Gleason score, secondary Gleason score, node status, and response to therapy using the RNA-Seq data from TCGA-PRAD cohort. Expression values are shown as $\log_2$ normalized count+1. Statistical significance was calculated using one-way ANOVA with Dunnett's multiple-comparisons posthoc test.

**B.** Violin plot showing association of *MALAT1* expression with biochemical recurrence in TCGA-PRAD cohort. *MALAT1* transcript values are shown as log*2* normalized count+1.

**C.** Box plots depicting *MALAT1* expression in multiple human cancer cell lines using RNA-Seq data retrieved from the CCLE database. Expression values are shown as FPKM.

**D.** Bar plots showing relative expression of *MALAT1* in prostate cancer cell line panel by quantitative PCR. The expression values are represented as $2^{-\delta}$CT.

**E.** Violin plot depicting expression of epithelial and mesenchymal markers in TCGA-PRAD cohort, stratified into "*MALAT1*-high" (n =125) and "*MALAT1*-low" (n = 125) groups based on the quartile-based normalization of *MALAT1* expression.

**F.** Bar plot depicting expression of *MALAT1* in 22RV1 and LNCaP cells transfected with sh*MALAT1* or shSCRM by quantitative PCR.

**G.** Same as E, except for the expression of stemness markers in the TCGA-PRAD cohort.

Experiments were performed with n = 3 biologically independent samples; data represents mean ± SEM. For panel F one-way ANOVA with Dunnett's multiple comparisons, posthoc test, and for panels, A,B,E and G two-tailed unpaired Student's t-test was applied. $*p \leq 0.05$ and $**p \leq 0.001$.

### Supplemental Figure 2

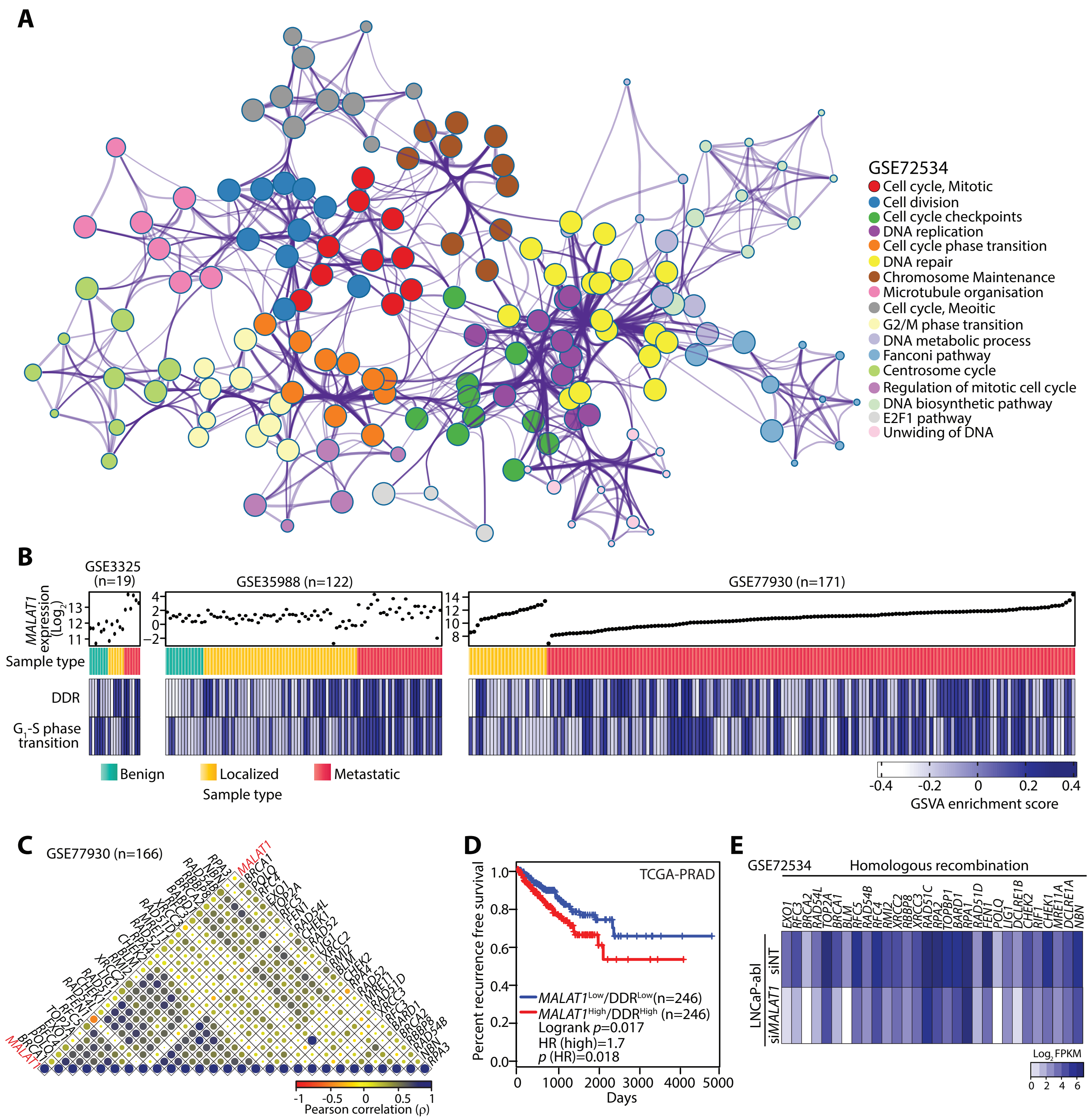


**Supplemental Figure 2: *MALAT1* regulates the DNA repair pathway in prostate cancer.**

**A.** Metascape enrichment network showing the downregulated pathways on silencing *MALAT1* in LNCaP-abl cells with a cutoff of *p*-value<0.01, minimum count>3, enrichment factor>1.5, and FDR>0.05. The network was visualized using Cytoscape, where each node represents an enriched term and its color indicates the cluster identity, while its size correlates with statistical significance. The nodes were clustered based on functional similarities wherein terms with a similarity score>0.3 were linked by an edge (the thickness of the edge represents the similarity score).

**B.** Heatmap depicting GSVA enrichment scores for DDR and $G_1$-S transition gene sets significantly enriched (FDR<0.05) in three prostate cancer patient cohorts, namely GSE35988, GSE77930, and GSE3325. The gene set enrichment scores values are expressed by the shades of blue and white. The topmost annotation in the heatmap indicates *MALAT1* expression and the second annotation represents the tumor subtype for the sample.

**C.** Correlogram representing Pearson correlation coefficient (ρ) between genes associated with DNA repair and *MALAT1* in prostate cancer patient samples from GSE77930 dataset (FDR adjusted, *p*<0.05). Correlation coefficients are expressed by shades of red and blue, and the size of dots is proportional to the strength of the correlation. Representative genes are marked on the sides of the correlogram.

**D.** Kaplan-Meier curve for relapse-free survival in TCGA-PRAD cohort categorized into two groups according to the median expression of *MALAT1* and DDR genes signature. The Red line represents high gene expression, whereas the blue line represents low gene expression. The *p*-values were computed by log-rank test.

**E.** Heatmap depicting the expression of differentially expressed DNA repair genes in the same cell as in A. Shades of blue represent log2 fold-change in gene expression.

### Supplemental Figure 3

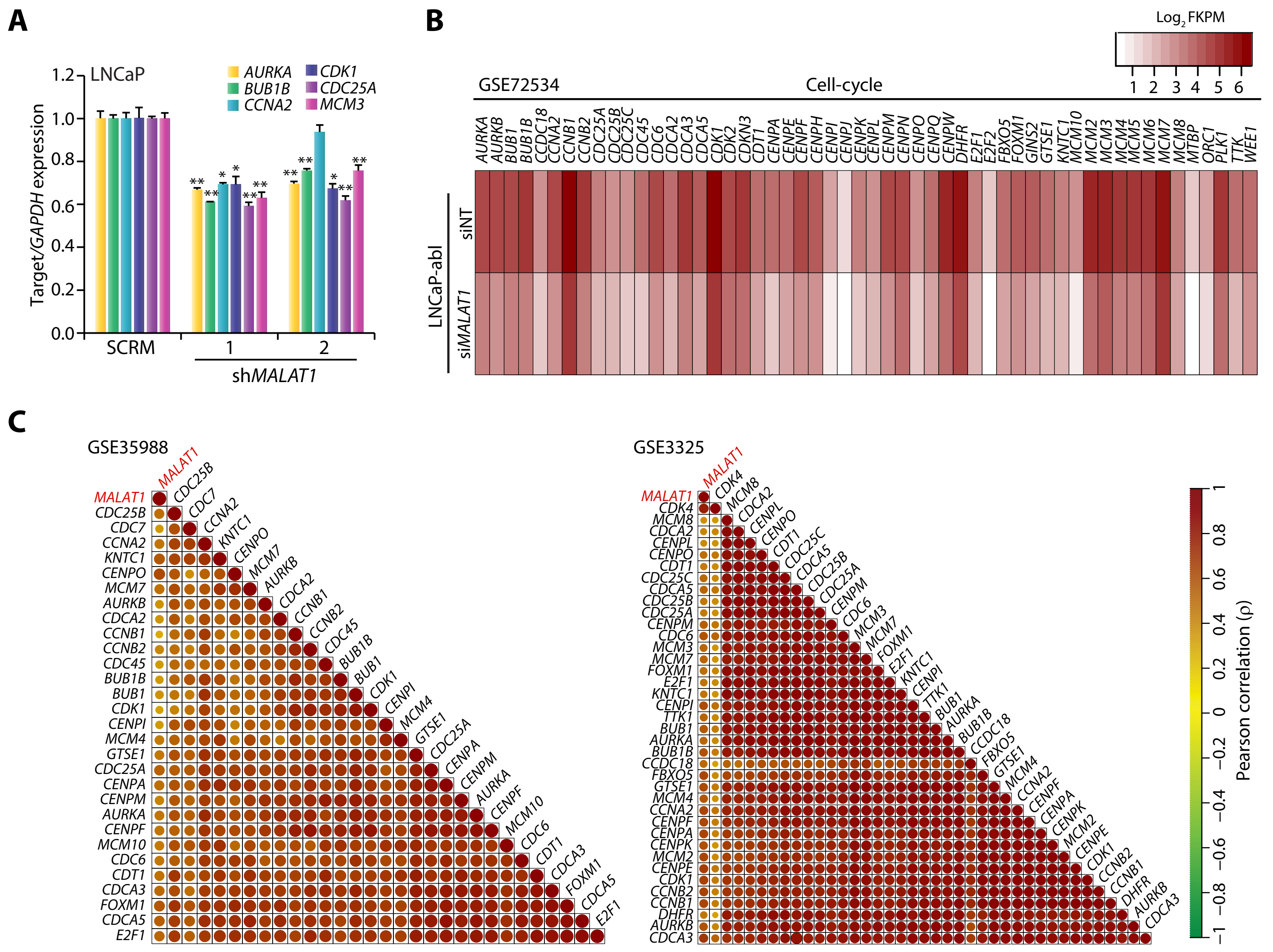


**Supplemental Figure 3: *MALAT1* depletion restrains the cell cycle in prostate cancer.**

**A.** QPCR depicting expression of cell-cycle markers in LNCaP-sh*MALAT1* and SCRM cells. Experiments was performed with n = 3 biologically independent samples; data represents mean ± SEM and significance was calculated using one –way ANOVA with Dunnett's multiple comparisons posthoc test. *$p \leq 0.05$ and **$p \leq 0.001$.

**B.** Heatmap depicting differential expression of genes associated with the cell cycle in LNCaP-abl-si*MALAT1* compared to LNCaP-abl-siCTL cells. Shades of red represent $\log_2$ fold-change in gene expression.

**C.** Correlogram representing Pearson correlation coefficient (ρ) between genes associated with cell cycle and *MALAT1* in prostate cancer patient samples from GSE35988 and GSE3325 dataset (FDR adjusted, p<0.05). Correlation coefficients are expressed by shades of green and red, and the size of dots is proportional to the strength of the correlation. Representative genes are marked on the sides of the correlogram.

### Supplemental Figure 4

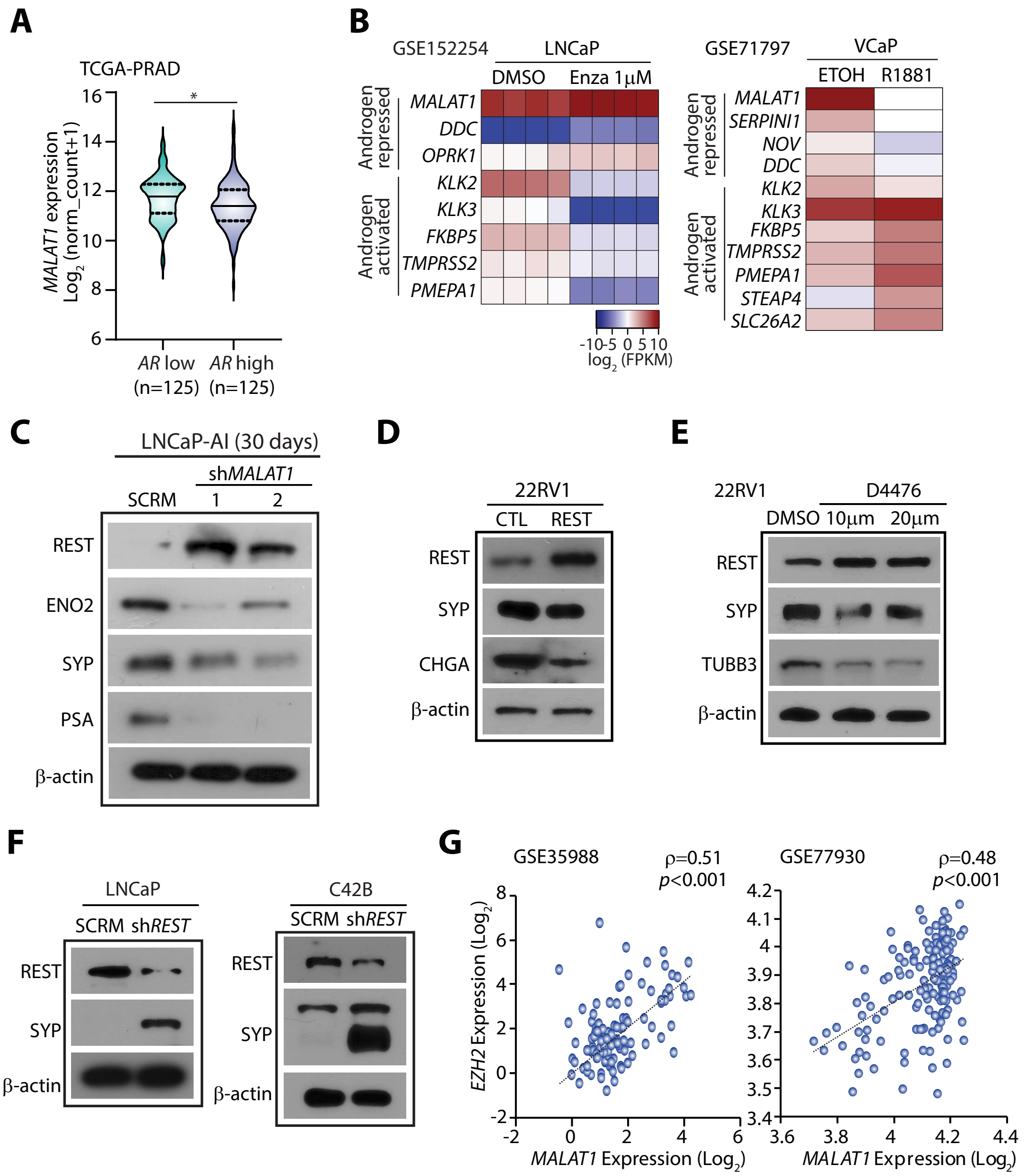


**Supplemental Figure 4: *MALAT1* modulates NE transdifferentiation by inhibiting REST.**

**A.** Violin plot depicting expression of *MALAT1* in TCGA-PRAD cohort, stratified into "*AR*-high" (n=125) and "*AR*-low" (n=125) groups based on the quartile-based normalization of *MALAT1* expression.

**B.** Heatmap depicting relative expression of androgen-regulated genes in LNCaP cells treated with enzalutamide (left, GSE152254) and androgen stimulated VCaP cells (right, GSE71797) for 24 hours. Shades of red and blue represent $\log_2$ fold-change in gene expression.

**C.** Immunoblot showing expression of NE markers and PSA in LNCaP-sh*MALAT1* and SCRM cells after 30 days of androgen deprivation.

**D.** Immunoblot showing expression of REST and NE markers in 22RV1 cells overexpressing REST and vector control. β-actin was used as an internal control.

**E.** Immunoblot showing expression of REST and NE markers in 22RV1 cells treated with the indicated concentrations of iCK1 (bottom panel). β-actin was used as an internal control.

**F.** Immunoblot depicting expression of REST and SYP in sh*REST* and SCRM LNCaP and C4-2B cells. β-actin was used as an internal control.

**G.** Scatter plot depicting the Pearson correlation between *MALAT1* and *EZH2* in prostate cancer patient datasets, namely GSE35988 and GSE77930. *MALAT1* and *EZH2* expression is reported as $\log_2$ median-centered ratio.
